# Characterization of the small *Arabidopsis thaliana* GTPase and ADP-ribosylation factor-like 2 protein TITAN 5

**DOI:** 10.1101/2023.04.27.538563

**Authors:** Inga Mohr, Amin Mirzaiebadizi, Sibaji K. Sanyal, Pichaporn Chuenban, Mohammad R. Ahmadian, Rumen Ivanov, Petra Bauer

## Abstract

Small GTPases function by conformational switching ability between GDP- and GTP-bound states in rapid cell signaling events. The ADP-ribosylation factor (ARF) family is involved in vesicle trafficking. Though evolutionarily well conserved, little is known about ARF and ARF-like GTPases in plants. Here, we characterized biochemical properties and cellular localization of the essential small ARF-like GTPase TITAN 5/HALLIMASCH/ARL2/ARLC1 (hereafter termed TTN5) from *Arabidopsis thaliana*. Two TTN5 variants were included in the study with point mutations at conserved residues, suspected to be functional for nucleotide exchange and GTP hydrolysis, TTN5^T30N^ and TTN5^Q70L^. We found that TTN5 had a very rapid intrinsic nucleotide exchange capacity with a conserved nucleotide switching mechanism. TTN5 acted as a non-classical small GTPase with a remarkably low GTP hydrolysis activity, suggesting it is likely present in GTP-loaded active form in the cell. We analyzed signals from yellow fluorescent protein (YFP)-tagged TTN5 and from *in situ* immunolocalization of hemagglutine-tagged HA_3_-TTN5 in Arabidopsis seedlings and in a transient expression system. Together with colocalization using endomembrane markers and pharmacological treatments the microscopic analysis suggests that TTN5 can be present at the plasma membrane and dynamically associated with membranes of vesicles, Golgi stacks and multivesicular bodies. While the TTN5^Q70L^ variant showed similar GTPase activities and localization behavior as wild-type TTN5, the TTN5^T30N^ mutant differed in some aspects.

Hence, the unusual capacity of rapid nucleotide exchange activity of TTN5 is linked with cell membrane dynamics, likely associated with vesicle transport pathways in the endomembrane system.

**Highlights:**

- The small ARF-like GTPase TTN5 has a very rapid intrinsic nucleotide exchange capacity with a conserved nucleotide switching mechanism
- Biochemical data classified TTN5 as a non-classical small GTPase, likely present in GTP-loaded active form in the cell
- YFP-TTN5 is dynamically associated with vesicle transport and different processes of the endomembrane system, requiring the active form of TTN5

## Introduction

A large variety of regulatory processes in signal transduction depends on guanine nucleotide-binding proteins of the GTPase family. Following the identification of common oncogenes (*HRAS, KRAS* and *NRAS*) a new class of GTPases has been recognized, that became known as the RAS superfamily of small GTPases (Bos 1988, Hall 1990, Kahn et al. 1992). RAS proteins have many conserved members in the eukaryotic kingdom. The RAS superfamily consists of five subfamilies in mammals: the Rat sarcoma (RAS), RAS homologs (RHO), RAS-like proteins in the brain (RAB), Ras-related nuclear proteins (RAN) and ADP-ribosylation factor (ARF) subfamilies (Kahn et al. 1992, Ahmadi et al. 2017). In *Arabidopsis thaliana* (Arabidopsis) only four families are represented, the ROP (Rho of plants), RAB, RAN and the ARF (Vernoud et al. 2003). The subfamilies are classified by sequence identity and characteristic sequence motifs with well-conserved regulatory functions within the cell (Kahn et al. 1992). Many mammalian small GTPases act as molecular switches in signal transduction. They switch from inactive GDP-loaded to active GTP-loaded GTPase form. The different activity states enable them to form differential complexes with proteins or act in tethering complexes to the target membrane. Small GTPases have usually low intrinsic GDP/GTP exchange and GTP hydrolysis activity and require the regulation by guanine nucleotide exchange factors (GEFs) and GTPase-activating proteins (GAPs). GEFs are potentially recruited to the inactive, GDP-bound GTPase at their site of action and accelerate GDP/GTP exchange leading to GTPase activation. GTP binding induces a conformational change of two regions referred to as switch I and II. The active, GTP-loaded GTPases exert their function *via* direct interaction with their effectors (Sztul et al. 2019, Nielsen 2020, Adarska et al. 2021) until their inactivation by GAPs, which stimulate the hydrolysis of GTP. Most of the known protein interactions important for their signaling functions occur in the active conformation of the GTPases. Members of the ARF family contain a conserved glycine at position 2 (Gly-2) for the characteristic *N*-myristoylation of an amphipathic helix (Kahn et al. 1992). ARF GTPases are often involved in vesicle-mediated endomembrane trafficking in mammalian cells and yeast (Just and Peränen 2016).

In plants, the activities of small GTPases and their functional environments in the plant cells are by far not understood well and only described in a very rudimentary manner. In particular, the ARF family of small GTPases is surprisingly poorly described in plants, although the Arabidopsis ARF family consists of twelve ARF, seven ARF-like and the associated SAR1 proteins (Singh et al. 2018). The best-studied plant ARF-GTPases, SAR1 and ARF1, act in the anterograde and retrograde vesicle transport between the endoplasmic reticulum (ER) and the Golgi. SAR1 is involved in COPII trafficking from the ER to the Golgi, whereas ARF1 participates in the opposite COPI pathway (Singh et al. 2018, Nielsen 2020). Another ARF-like protein, ARL1, has perhaps a role in endosome-to-Golgi trafficking (Latijnhouwers et al. 2005, Stefano et al. 2006). These roles of ARF1 and SAR1 in COPI and II vesicle formation within the endomembrane system are well conserved in eukaryotes which raises the question of whether other plant ARF members are also involved in functioning of the endomembrane system. A recent study showed Golgi-related localization for some ARF and ARF-like proteins (Niu et al. 2022) promoting a general involvement of the ARF family in the endomembrane system.

TITAN 5 (TTN5)/HALLIMASCH (HAL)/ARF-LIKE 2 (ARL2), ARLC1, from here on referred to as TTN5, is essential in plant development. It was identified in two independent screens for abnormal embryo mutants. The *ttn5* loss-of-function mutants are arrested soon after cell division of the fertilized egg cell, indicating a fundamental, potentially housekeeping role in cellular activities (Mayer et al. 1999, McElver et al. 2000, Lloyd and Meinke 2012). TTN5 is closely related in sequence to human ADP-ribosylation factor-like 2 (HsARL2). HsARL2 has high nucleotide dissociation rates, being up to 4000-fold faster compared to RAS (Hanzal-Bayer et al. 2005, Veltel et al. 2008). HsARL2 is associated with different functions in cells, ranging from microtubule development, also identified for yeast and Caenorhabditis homologs (Bhamidipati et al. 2000, Fleming et al. 2000, Radcliffe et al. 2000, Antoshechkin and Han 2002, Tzafrir et al. 2002, Mori and Toda 2013), adenine nucleotide transport in mitochondria (Sharer et al. 2002) and control of phosphodiesterase activity in cilia (Ismail et al. 2011, Fansa and Wittinghofer 2016). It is thought that HsARL2 requires fast-acting nucleotide exchange and participates in different cellular processes that depend on developmental and tissue-specific factors. With regard to the plant ortholog, it is still completely unknown, which cellular roles TTN5 fulfills in plants. Until today, the function of this small plant GTPase remains elusive at the molecular level. Besides lacking knowledge of the physiological context, the GTPase characteristics and properties of the TTN5 enzyme are not yet demonstrated.

Here, we show by stopped-flow fluorimetry kinetic assays that TTN5 is a functional small GTPase with conserved GTP hydrolysis and very fast nucleotide exchange characteristics. Based on fluorescence microscopy combined with pharmacological treatments, TTN5 may be located at the plasma membrane and within the endomembrane system. Our study enables future investigation of the cellular-physiological functions of this small GTPase.

## Results

### TTN5 exhibited atypical characteristics of rapid nucleotide exchange and slow GTP hydrolysis

There is a higher sequence similarity of TTN5 with its animal ARL2 ortholog than to any Arabidopsis ARF/ARL proteins (Figure 1A) (McElver et al. 2000, Vernoud et al. 2003). Several observations indicate that TTN5 plays a fundamental and essential role in cellular activities. Loss of function of TTN5 causes a very early embryo-arrest phenotype (Mayer et al. 1999, McElver et al. 2000). An essential TTN5 function is also reflected by its regulation and ubiquitous gene expression during plant development and in the root epidermis revealed in public RNA-seq data sets of organ and single cell analysis of roots (Supplementary Figure S1A, B). *TTN5* is strongly expressed during early embryo development where cell division, cell elongation and cell differentiation take place (Supplementary Figure S1C). Hence, *TTN5* is expressed and presumably functional in very fundamental processes in cells with stronger expression when cells grow and divide.

**Figure 1:**
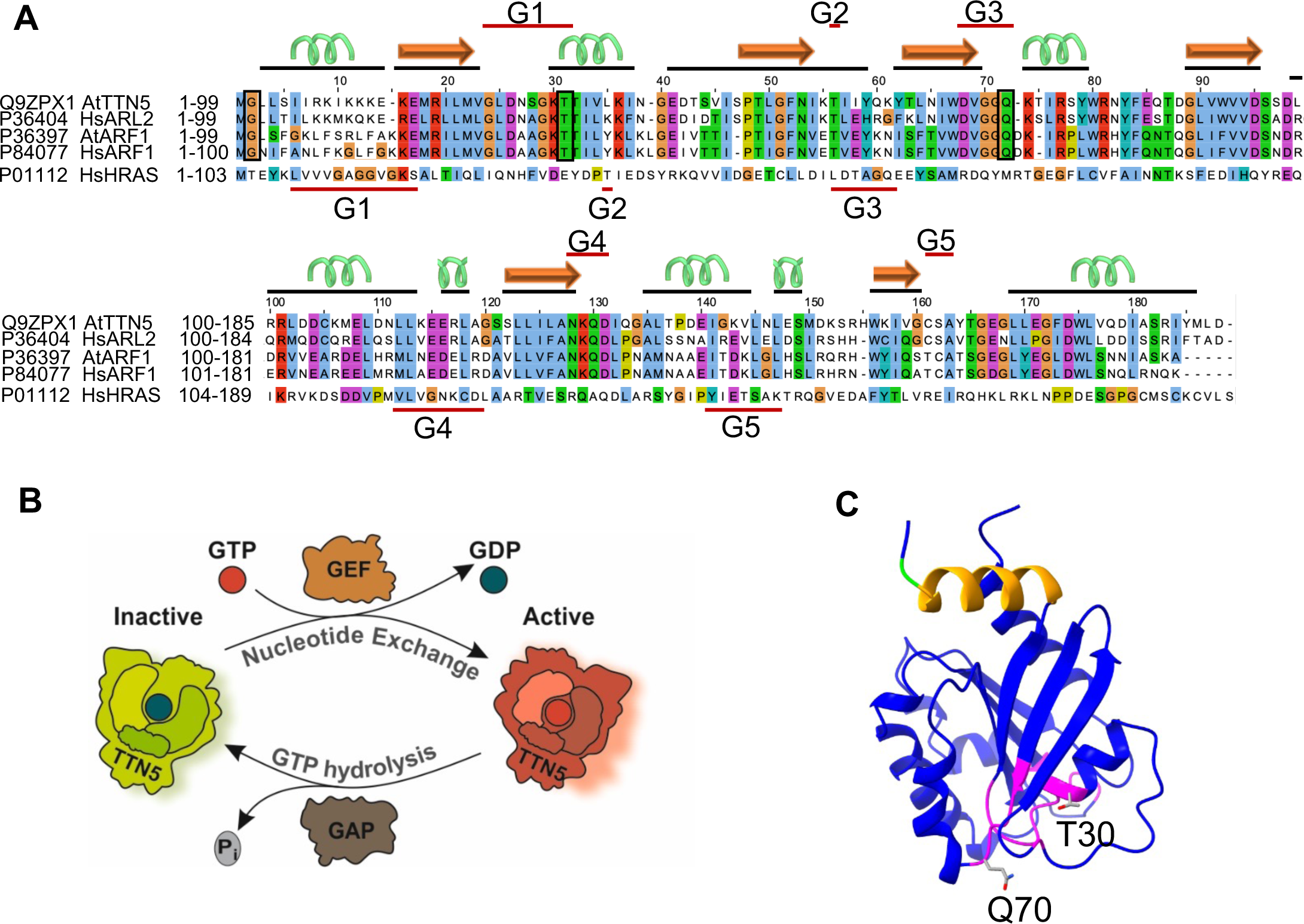
TTN5 is predicted to be a functional small ARF-like GTPase with nucleotide exchange capacity. (A), Sequence alignment of TTN5 with its human homolog ARL2, Arabidopsis, human ARF1 and human HRAS created with Jalview (Waterhouse et al. 2009). The conserved G-motifs (G1-G5; indicated by red lines) are defined for the TTN5 and HRAS sequence. The secondary structure of TTN5 is depicted by black lines and corresponding cartoon (α-helix in green; β-sheet in orange). Here mentioned conserved residues in ARF/ARL proteins are highlighted by boxes; Gly-2, and mutated Thr-30 and Gln-70. TTN5^T30N^ is expected to have a low nucleotide exchange capacity, while TTN5^Q70L^ is expected to have a low GTPase hydrolysis activity. (B), Model of the predicted GTPase nucleotide exchange and hydrolysis cycle of TTN5. TTN5 switches from an inactive GDP-loaded form to an active GTP-loaded one. GDP to GTP nucleotide exchange and GTP hydrolysis may be aided by a guanidine exchange factor (GEF) and a GTPase-activating protein (GAP). (C), Predicted protein structural model of TTN5; magenta, marks the GTP-binding pocket; N-terminal amphipathic helix is highlighted in orange; conserved Gly-2 in green; T30 and Q70, mutagenized in this study, shown in sticks. The model was generated with AlphaFold (Jumper et al. 2021), and adaptation was done with UCSF ChimeraX 1.2.5 (Goddard et al. 2018).

The molecular switch functions can be presumed for TTN5 based on its sequence similarity and structural prediction (Figure 1B, C). HsARL2 has a fast GDP/GTP exchange characteristic (Hanzal-Bayer et al. 2005, Veltel et al. 2008). However, it had not been known whether the plant TTN5 has similar or different GTPase characteristics as its animal counterparts. In this study, we characterized the nucleotide binding and GTP hydrolysis properties of TTN5^WT^ and two of its mutants, TTN5^T30N^ and TTN5^Q70L^, using heterologously expressed and purified proteins and *in vitro* biochemical assays, as previously established for human GTPases (Eberth and Ahmadian 2009). The experimental workflow is illustrated in Supplementary Figure S2A-E. The dominant-negative TTN5^T30N^ is assumed to preferentially bind GEFs, sequestering them from their proper context, while the constitutively active TTN5^Q70L^ is thought to be defective in hydrolyzing GTP. Equivalent mutants have been frequently used and characterized in previous studies (Scheffzek et al. 1997, Zhou et al. 2006, Newman et al. 2014). We monitored the real-time kinetics of the interactions of fluorescent guanine nucleotides using stopped-flow fluorimetry suited for very rapid enzymatic reactions (Figure 2A-C). 2-deoxy-3-O-N-methylanthraniloyl-deoxy-GDP (mdGDP) and GppNHp (mGppNHp), a non-hydrolyzable GTP analog, were used to mimic GDP and GTP binding to TTN5 proteins. This approach allowed us to monitor real-time kinetics and quantify nucleotide association and dissociation characteristics of small GTPases, such as HsARL2 and HsARL3 (Hillig et al. 2000, Hanzal-Bayer et al. 2005, Veltel et al. 2008, Zhang et al. 2018). The kinetics allow us to determine the association rate constant (k_on_) and the dissociation rate constant (k_off_), respectively. The k_on_ value is defined as the rate of nucleotide binding to the GTPase to form the GTPase-nucleotide complex (Figure 2B) whereas the k_off_ value describes the rate of nucleotide dissociation from the GTPase (Figure 2C). We found that TTN5 proteins were able to bind both nucleotides, with the exception of mGppNHp binding by TTN5^T30N^ (Supplementary Figures S3A-F, S4A-E). TTN5^Q70L^ revealed the highest k_on_ value for mGDP binding (0.401 µM^-1^s^-1^), which was 9-fold higher compared to TTN5^WT^ (0.044 µM^-1^s^-1^) and TTN5^T30N^ (0.048 µM^-1^s^-1^), respectively (Figure 2D; Supplementary Figure S3D-F). The k_on_ values for mGppNHp binding were 2-fold lower for TTN5^WT^ (0.029 µM^-1^s^-1^) and TTN5^Q70L^ (0.222 µM^-1^s^-1^) compared to those for mGDP binding, respectively (Figure 2E; Supplementary Figure S4C, D). The differences in k_on_ for the respective nucleotide binding were small. However, TTN5^Q70L^ showed a 7.5-fold faster mGppNHp binding than TTN5^WT^. A remarkable observation was that we were not able to monitor the kinetics of mGppNHp association with TTN5^T30N^ but observed its dissociation (k_off_ = 0.026 s^-1^; Figure 2E). To confirm the binding capability of TTN5^T30N^ with mGppNHp, we measured the mGppNHp fluorescence in real-time before and after titration of nucleotide-free TTN5^T30N^. As shown in Supplementary Figure S4B, the binding of mGppNHp to TTN5^T30N^ occurs so fast that it was not possible to resolve the rate of association.

**Figure 2:**
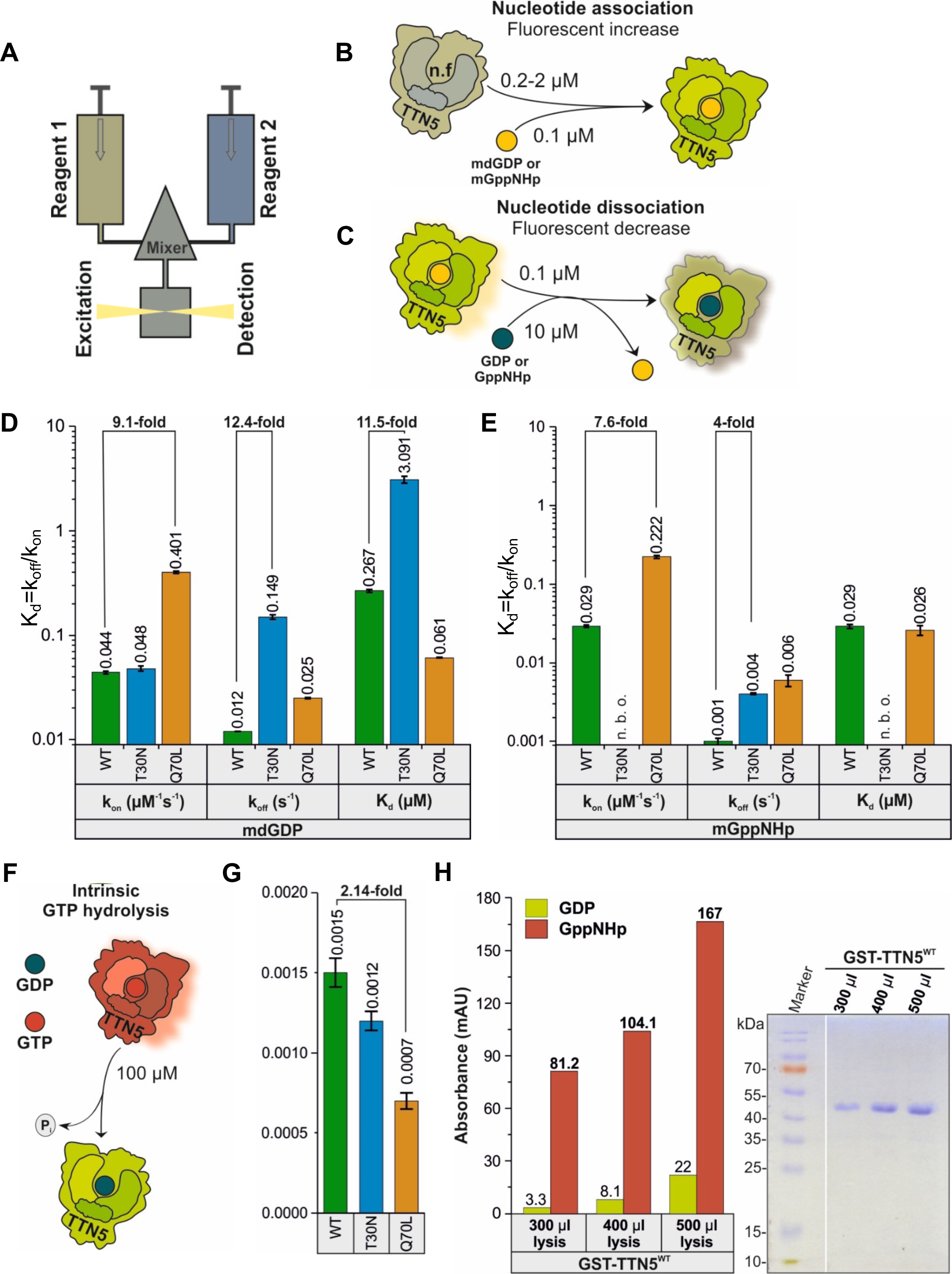
Biochemical properties of TTN5 proteins suggest that TTN5 is present in a GTP-loaded active form in cells. (A), Schematic illustration of the stopped-flow fluorescence device for monitoring the nucleotide-binding kinetics of the purified TTN5 protein heterologously expressed in bacteria (Supplementary Figure S2A-D). It consists of two motorized, thermostated syringes, a mixing chamber and a fluorescence detector. Two different reagents 1 and 2 are rapidly mixed and transferred to a fluorescence detection cell within 4 ms. One of the reagents must contain a fluorescent reporter group. Here, mdGDP and mGppNHp were used to mimic GDP and GTP. (B), Schematic illustration of the nucleotide association. Nucleotide-free TTN5 (reagent 1; preparation see Supplementary Figure S2E) was rapidly mixed with mdGDP (reagent 2). A fluorescence increase is expected upon association of mdGDP with TTN5. Similar measurements are performed with mGppNHp instead of mdGDP. (C), Schematic illustration of the intrinsic nucleotide dissociation. mdGDP-bound TTN5 (reagent 1) is mixed with a molar excess of GDP (reagent 2). A fluorescence decrease is expected upon mdGDP dissociation from TTN5 and binding of free GDP. Similar measurements are performed with mGppNHp. (D-E), Kinetics of association and dissociation of fluorescent nucleotides mdGDP (D) or mGppNHp (E) with TTN5 proteins (WT, TTN5^T30N^, TTN5^Q70L^) are illustrated as bar charts. The association of mdGDP (0.1 µM) or mGppNHp (0.1 µM) with increasing concentration of TTN5^WT^, TTN5^T30N^ and TTN5^Q70L^ was measured using a stopped-flow device (see A, B; data see Supplementary Figure S3A-F, S4A-E). Association rate constants (k_on_ in µM^-1^s^-1^) were determined from the plot of increasing observed rate constants (k_obs_ in s^-1^) against the corresponding concentrations of the TTN5 proteins. Intrinsic dissociation rates (k_off_ in s^-1^) were determined by rapidly mixing 0.1 µM mdGDP-bound or mGppNHp-bound TTN5 proteins with the excess amount of unlabeled GDP (see A, C, data see Supplementary Figure S3G-I, S4F-H). The nucleotide affinity (dissociation constant or K_d_ in µM) of the corresponding TTN5 proteins was calculated by dividing k_off_ by k_on_. When mixing mGppNHp with nucleotide-free TTN5^T30N^, no binding was observed (n.b.o.) under these experimental conditions. (F-G), GTP hydrolysis of TTN5 proteins determined by HPLC. (F), Schematic illustration of the GTP hydrolysis measurement. (G), GTP-bound TTN5 proteins (100 µM) were incubated at room temperature at different time points before injecting them on a reversed-phase HPLC system. Evaluated data (data see Supplementary Figure S5) resulted in the determination of the GTP hydrolysis rates (k_cat_ in s^-1^) illustrated as bar charts. (H), TTN5 accumulated in a GTP-loaded active form. GST-TTN5^WT^ (46.5 kDa) was purified from bacterial cell lysates at three different volumes in the presence of 0.1 µM unbound free GppNHp using glutathione beads. The nucleotide contents and the protein purities were determined by HPLC and Coomassie Blue-stained SDS-polyacrylamide gel electrophoresis. The presence of much higher amounts of GppNHp-bound versus GDP-bound GST-TTN5 protein indicates that TTN5 rapidly exchanged bound nucleotide und accumulated in this state.

We next measured the dissociation (k_off_) of mdGDP and mGppNHp from the TTN5 proteins in the presence of excess amounts of GDP and GppNHp, respectively (Figure 2C) and found interesting differences (Figure 2D, E; Supplementary Figures S3G-I, S4F-H). First, TTN5^WT^ showed a k_off_ value (0.012 s^-1^ for mGDP) (Figure 2D; Supplementary Figure S3G), which was 100-fold faster than those obtained for classical small GTPases, including RAC1 (Haeusler et al. 2006) and HRAS (Gremer et al. 2011), but very similar to the k_off_ value of HsARF3 (Fasano et al. 2022). Second, the k_off_ values for mGDP and mGppNHp, respectively, were in a similar range between TTN5^WT^ (0.012 s^-1^ mGDP and 0.001 s^-1^ mGppNHp) and TTN5^Q70L^ (0.025 s^-1^ mGDP and 0.006 s^-1^ mGppNHp), respectively, but the k_off_ values differed 10-fold between the two nucleotides mGDP and mGppNHp in TTN5^WT^ (k_off_ = 0.012 s^-1^ versus k_off_ = 0.001 s^-1^; Figure 2D, E; Supplementary Figure S3G, I, S4F, H). Thus, mGDP dissociated from proteins 10-fold faster than mGppNHp. Third, the mGDP dissociation from TTN5^T30N^ (k_off_ = 0.149 s^-1^) was 12.5-fold faster than that of TTN5^WT^ and 37-fold faster than the mGppNHp dissociation of TTN5^T30N^ (k_off_ = 0.004 s^-1^) (Figure 2D, E; Supplementary Figure S3H, S4G). Mutants of CDC42, RAC1, RHOA, ARF6, RAD, GEM and RAS GTPases, equivalent to TTN5^T30N^, display decreased nucleotide binding affinity and therefore tend to remain in a nucleotide-free state in a complex with their cognate GEFs (Erickson et al. 1997, Ghosh et al. 1999, Radhakrishna et al. 1999, Jung and Rösner 2002, Kuemmerle and Zhou 2002, Wittmann et al. 2003, Nassar et al. 2010, Huang et al. 2013, Chang and Colecraft 2015, Fisher et al. 2020, Shirazi et al. 2020). Since TTN5^T30N^ exhibits fast guanine nucleotide dissociation, these results suggest that TTN5^T30N^ may also act in either a dominant-negative or fast-cycling manner as reported for other GTPase mutants (Fiegen et al. 2004, Wang et al. 2005, Fidyk et al. 2006, Klein et al. 2006, Soh and Low 2008, Sugawara et al. 2019, Aspenström 2020).

The dissociation constant (K_d_) is calculated from the ratio k_off_/k_on_, which inversely indicates the affinity of the interaction between proteins and nucleotides (the higher K_d_, the lower affinity). Interestingly, TTN5^WT^ binds mGppNHp (K_d_ = 0.029 µM) 10-fold tighter than mGDP (K_d_ = 0.267 µM), a difference, which was not observed for TTN5^Q70L^ (K_d_ for mGppNHp = 0.026 µM, K_d_ for mGDP = 0.061 µM) (Figure 2D, E). The lower affinity of TTN5^WT^ for mdGDP compared to mGppNHp brings us one step closer to the hypothesis that classifies TTN5 as a non-classical GTPase with a tendency to accumulate in the active (GTP-bound) state (Jaiswal et al. 2013). The K_d_ value for the mGDP interaction with TTN5^T30N^ was 11.5-fold higher (3.091 µM) than for TTN5^WT^, suggesting that this mutant exhibited faster nucleotide exchange and lower affinity for nucleotides than TTN5^WT^. Similar as other GTPases with a T30N exchange, TTN5^T30N^ may behave in a dominant-negative manner in signal transduction (Vanoni et al. 1999).

To get hints on the functionalities of TTN5 during the complete GTPase cycle, it was crucial to determine its ability to hydrolyze GTP. Accordingly, the catalytic rate of the intrinsic GTP hydrolysis reaction, defined as k_cat_, was determined by incubating 100 µM GTP-bound TTN5 proteins at 25°C and analyzing the samples at various time points using a reversed-phase HPLC column (Figure 2F; Supplementary Figure S5). The determined k_cat_ values were quite remarkable in two respects (Figure 2G). First, all three TTN5 proteins, TTN5^WT^, TTN5^T30N^ and TTN5^Q70L^, showed quite similar k_cat_ values (0.0015 s^-1^, 0.0012 s^-1^, 0.0007 s^-1^; Figure 2G; Supplementary Figure S5). The GTP hydrolysis activity of TTN5^Q70L^ was quite high (0.0007 s^-1^). This was unexpected because, as with most other GTPases, the glutamine mutations at the corresponding position drastic impair hydrolysis, resulting in a constitutively active GTPase in cells (Hodge et al. 2020, Matsumoto et al. 2021). Second, the k_cat_ value of TTN5^WT^ (0.0015 s^-1^) although quite low as compared to other GTPases (Jian et al. 2012, Esposito et al. 2019), was 8-fold lower than the determined k_off_ value for mGDP dissociation (0.012 s^-1^) (Figure 2E). This means that a fast intrinsic GDP/GTP exchange versus a slow GTP hydrolysis can have drastic effects on TTN5 activity in resting cells, since TTN5 can accumulate in its GTP-bound form, unlike the classical GTPase (Jaiswal et al. 2013). To investigate this scenario, we pulled down GST-TTN5 protein from bacterial lysates in the presence of an excess amount of GppNHp in the buffer using glutathione beads and measured the nucleotide-bound form of GST-TTN5 using HPLC. As shown in Figure 2H, isolated GST-TTN5 increasingly bonds GppNHp, indicating that the bound nucleotide is rapidly exchanged for free nucleotide (in this case GppNHp). This is not the case for classical GTPases, which remain in their inactive GDP-bound forms under the same experimental conditions (Walsh et al. 2019, Hodge et al. 2020).

In summary, the TTN5 sequence not only contains conserved regions necessary for nucleotide binding but theTTN5 protein also binds nucleotides detectably. Interestingly, the slow intrinsic GTP hydrolysis rates in combination with the high dissociation rates for GDP suggest that TTN5 tends to exist in a GTP-loaded form. A fast intrinsic GDP/GTP exchange and a slow GTP hydrolysis can have drastic effects on TTN5 activity in cells under resting/unstimulated conditions, as TTN5 can accumulate in its GTP-bound form, unlike the classical GTPases (Jaiswal et al. 2013). On the other hand, the originally suspected constitutively active TTN5^Q70L^ still has intrinsic GTPase activity, while the T30N variant exhibits a low affinity for mGDP. Therefore, we propose that TTN5 exhibits the typical functions of a small GTPase based on *in vitro* biochemical activity studies, including guanine nucleotide association and dissociation, but emphasizes its divergence among the ARF GTPases by its kinetics.

### TTN5 may be a highly dynamic protein and localize to different intracellular compartments

Several ARF GTPases function in vesicle transport and are located at various membranous sites linked with the endomembrane compartments in eukaryotes (Vernoud et al. 2003). Localization had not been comprehensively studied for TTN5. To obtain hints where in a cell TTN5 may be localized, we first created transgenic Arabidopsis lines constitutively expressing YFP-tagged TTN5 (pro35S::YFP-TTN5) and its two mutant forms (pro35S::YFP-TTN5^T30N^, pro35S::YFP-TTN5^Q70L^) and investigated the localization in 6-day-old seedlings in the epidermis of cotyledons, hypocotyls, root hair zone and in root tips (Figure 3A; Supplementary Figure S6A). The microscopic observations were made in different planes of the tissues, e.g. inside the cells across the vacuoles (Supplementary Figure S6) and underneath the plasma membrane at the cell peripheries (Figure 3). We chose the investigation of YFP-TTN5 in the epidermis as this is a tissue where *TTN5* transcripts were detected in plants (Supplementary Figure S1B). YFP signals in YFP-TTN5 seedlings were detected in the nucleus, in the cytoplasm and at or in close proximity to the plasma membrane in the epidermal cotyledon cells (Supplementary Figure S6B). The same localization patterns were found for mutant YFP-TTN5 signals (Supplementary Figure S6C-D). The YFP-signals in YFP-TTN5, YFP-TTN5^T30N^ and YFP-TTN5^Q70L^ seedlings were also present in a similar pattern in the stomata (Figure 3B-D). In hypocotyls of seedlings, intracellular YFP signal was observed in nuclei and in close proximity to or at the plasma membrane with all three YFP-TTN5 forms (Supplementary Figure S6E-G). Investigation of the root hair zone showed YFP signals in the cytoplasm and at the plasma membrane of root hairs (Supplementary Figure S6H-J). In the root tip, YFP signal was detectable inside the cytoplasm and in nuclei (Supplementary Figure S6K). The pattern was similar for YFP-TTN5^T30N^ and YFP-TTN5^Q70L^ (Supplementary Figure S6L-M). Fluorescent signal in YFP-TTN5, YFP-TTN5^T30N^ and YFP-TTN5^Q70L^ seedlings inside the cytoplasm was confined to punctate structures indicating that the signals were present in cytosolic vesicle-like structures together with free signals in the cytosol. This localization pattern was also present in leaf epidermal cells and stomata of the cotyledons (Figure 3B-D), in the hypocotyls (Figure 3E-G) and in the cells of the root hair zones and in the root hairs (Figure 3H-J). These observed structures point to an association of TTN5 with vesicle and endomembrane trafficking. A closer inspection of the dynamics of these structures in the leaf epidermis of cotyledons showed high mobility of fluorescent signal within the cells (Supplementary Video Material S1A-C), likewise in hypocotyl cells (Supplementary Video Material S1D). Interestingly, the mobility of these punctate structures differed within the cells when the mutant YFP-TTN5^T30N^ was observed in hypocotyl epidermis cells, but not in the leaf epidermis cells (Supplementary Video Material S1E, compare with S1B) nor was it the case for the YFP-TTN5^Q70L^ mutant (Supplementary Video Material S1F, compare with S1E). We detected approximately half of the cells within the hypocotyl epidermis with slowed-down or completely arrested movement for YFP-TTN5^T30N^, in contrast to YFP-TTN5 and YFP-TTN5^Q70L^ (Supplementary Video Material S1D-F). This loss of fluorescence signal mobility in YFP-TTN5^T30N^ seedlings may be a consequence of missing effector interaction. We did not observe the blocked mobility for fluorescent signals in cells expressing YFP-TTN5, YFP-TTN5^T30N^ and YFP-TTN5^Q70L^ in the root elongation zone (Supplementary Video Material S1G-I). No mobility of YFP fluorescence signal was visible in root tip cells for any YFP-TTN5 form (Supplementary Video Material S1J-L).

**Figure 3:**
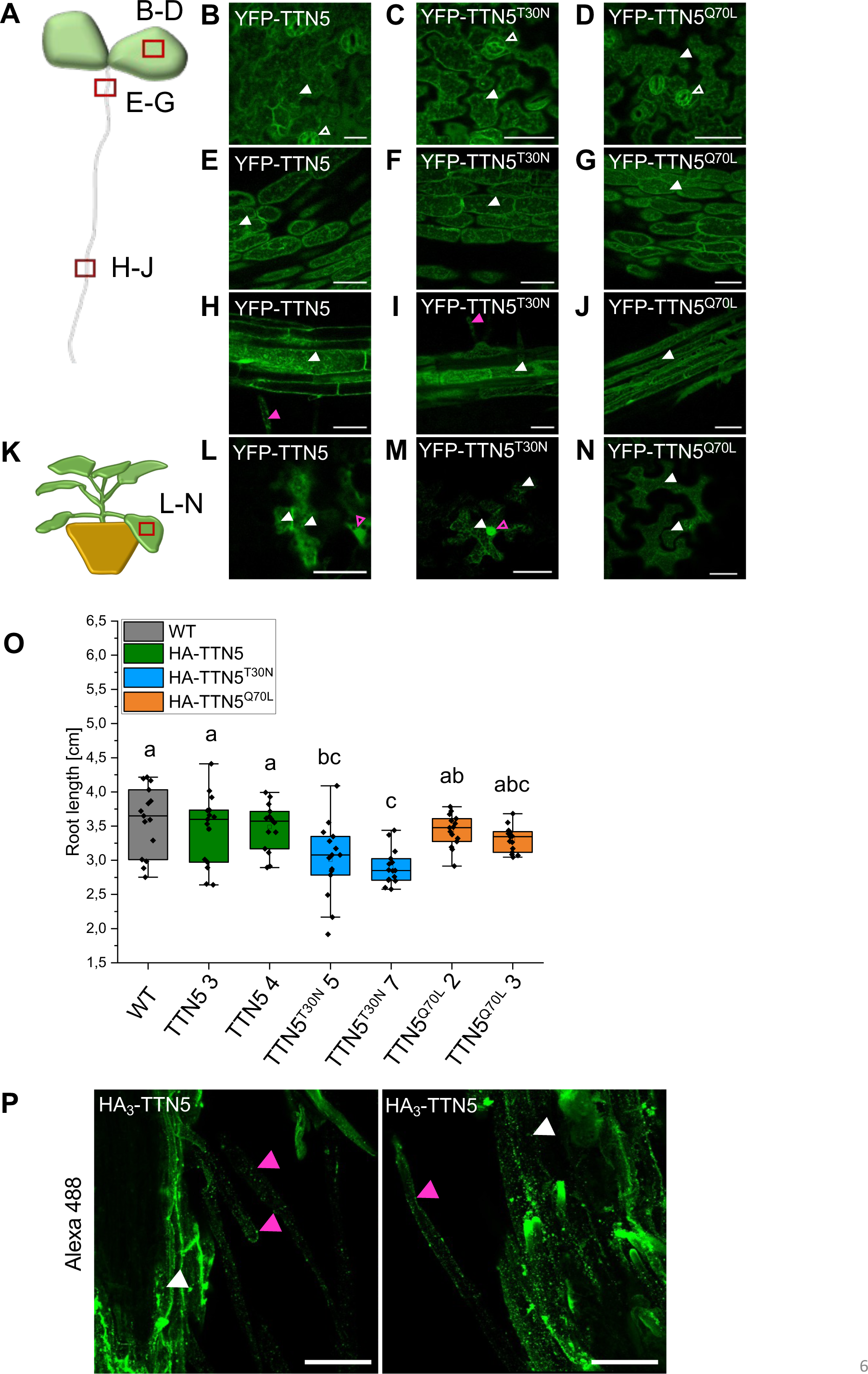
TTN5 may be present in punctate structures in seedlings. Microscopic observations of YFP fluorescence signals were made in a plane underneath the plasma membrane at the cell peripheries. (A), Schematic representation of an Arabidopsis seedling. Images were taken at three different positions of the seedlings and imaged areas are indicated by a red rectangle. (B-J), Analysis of YFP-TTN5, YFP-TTN5^T30N^ and YFP-TTN5^Q70L^ Arabidopsis seedlings *via* fluorescent confocal microscopy. (B-D), Fluorescence signals observed in stomata (indicated by empty white arrowhead) and in the epidermis of cotyledons in punctate structures (indicated by filled white arrowhead). (E-G), Localization in the hypocotyls showed the same pattern of punctate structures. (H-J), Signals were present in punctate structures in the root hair zone and in root hairs (indicated by filled magenta arrowhead). (K), Schematic representation of a *N. benthamiana* plant, used for leaf infiltration for transient expression. Imaged area is indicated by a red rectangle. (L-N), YFP fluorescence signals in *N. benthamiana* leaf epidermal cells expressing YFP-TTN5, YFP-TTN5^T30N^ and YFP-TTN5^Q70L^. Signals were present in punctate structures (indicated by white arrowheads) and in the nucleus (indicated by empty magenta arrowheads). Scale bar 50 μm. (O), Root length measurement of HA_3_-TTN5, HA_3_-TTN5^T30N^ and HA_3_-TTN5^Q70L^ Arabidopsis lines in comparison with non-transgenic wild type (WT). Seedlings were grown for 10 days on Hoagland plates. Only HA_3_-TTN5^T30N^ showed a slightly reduced root length compared to WT, whereas HA_3_-TTN5 and HA_3_-TTN5^Q70L^ did not have a divergent phenotype. Analysis was conducted in replicates (n = 14). One-way ANOVA with Tukey post-hoc test was performed. Different letters indicate statistical significance (p < 0.05). (P), Two representative images of whole-mount immunostaining of HA_3_-TTN5 seedling roots in the root differentiation zone (rabbit α-HA primary antibody, Alexa-488-labeled secondary α-rabbit antibody). Alexa-488 signals were present in punctate structures in root cells (indicated by filled white arrowhead) and in root hairs (indicated by filled magenta arrowhead) comparable to YFP signals (Figure 3H-J). Images are presented in a maximum intensity projection of several z-layers for a better visualization. Scale bar 50 µm.

To evaluate the Arabidopsis data and to better visualize YFP-TTN5, we expressed YFP-TTN5 constructs transiently in *Nicotiana benthamiana* leaf epidermis cells. We found that fluorescent signals in YFP-TTN5-, YFP-TTN5^T30N^- and YFP-TTN5^Q70L^-expressing cells were also all localized at or in close proximity to the plasma membrane and in several cytosolic punctate structures, apart from the nucleus, similar to Arabidopsis cotyledons, hypocotyls and root hair zones (Figure 3K-N; Supplementary Figure S6N-Q). Additionally, YFP signals were also detected in a net-like pattern typical for ER localization (Figure 3M, N). This showed that localization of fluorescent signal was similar between Arabidopsis epidermis cells and *N. benthamiana* leaf epidermis.

It should be noted that the 35S promoter-driven YFP-TTN5 constructs did not complement the embryo-lethal phenotype of *ttn5-1* (Supplementary Figure S7A, B). We also found multiple YFP bands in α-GFP Western blot analysis using YFP-TTN5 Arabidopsis seedlings. Besides the expected and strong 48 kDa YFP-TTN5 band, we observed three weak bands ranging between 26 to 35 kDa (Supplementary Figure S7C). We cannot explain the presence of these small protein bands. They might correspond to free YFP, to proteolytic products or potentially to proteins produced from aberrant transcripts with perhaps alternative translation start or stop sites. On the other side, a triple hemagglutinin-tagged HA_3_-TTN5 driven by the 35S promoter did complement the embryo-lethal phenotype of *ttn5-1* (Supplementary Figure S7D, E). α-HA Western blot control performed with plant material from HA_3_-TTN5 seedlings showed a single band at the correct size, but no band that was 13 to 18 kDa smaller (Supplementary Figure S7D). Hence, the inability of YFP-TTN5 to complement the embryo-lethal phenotype was presumably due to the YFP-tag which was rather large compared with the small GTPase and larger than the relatively small HA_3_-tag. Interestingly, HA_3_-TTN5^T30N^ seedlings presented a root length phenotype, whereas HA_3_-TTN5 and HA_3_-TTN5^Q70L^ seedlings had no obvious phenotype compared to wild type plants. HA_3_-TTN5^T30N^ roots were shorter than those of HA_3_-TTN5^Q70L^ and HA_3_-TTN5, which can be due to the atypical biochemical TTN5^T30N^ characteristics (Figure 3O).

To verify that the localization patterns observed with the YFP-TTN5 constructs are representative of a functional TTN5, we performed immunofluorescence staining against the HA_3_-tag in roots of HA_3_-TTN5 seedlings and compared the localization patterns (Figure 3P). Alexa 488-labeled α-HA antibody staining reflected HA_3_-TTN5 localization and signals were visible in root cells and root hairs as expected. Signals were mostly present in punctate structures close to the plasma membrane and in the cytosol (Figure 3P), fitting with above described fluorescence signals obtained with the YFP-TTN5 plants.

For a more detailed investigation of HA_3_-TTN5 subcellular localization, we then performed co-immunofluorescence staining with an Alexa 488-labeled antibody recognizing the Golgi and TGN marker ARF1, while detecting HA_3_-TTN5 with an Alexa 555-labeled antibody (Robinson et al. 2011, Singh et al. 2018) (Figure 4A). ARF1-Alexa 488 staining was clearly visible in punctate structures representing presumably Golgi stacks (Figure 4A, Alexa 488), as previously reported (Singh et al. 2018). Similar structures were obtained for HA_3_-TTN5-Alexa 555 staining (Figure 4A, Alexa 555). But surprisingly, colocalization analysis demonstrated that the HA_3_-TTN5-labeled structures were mostly not colocalizing and thus distinct from the ARF1-labeled ones (Figure 4A). Yet the HA_3_-TTN5- and ARF1-labeled structures were in close proximity to each other (Figure 4A). We hypothesized that the HA_3_-TTN5 structures can be connected to intracellular trafficking steps. To test this, we performed brefeldin A (BFA) treatment, a commonly used tool in cell biology for preventing dynamic membrane trafficking events and vesicle transport involving the Golgi. BFA is a fungal macrocyclic lactone that leads to a loss of *cis*-cisternae and accumulation of Golgi stacks, known as BFA-induced compartments, up to the fusion of the Golgi with the ER (Ritzenthaler et al. 2002, Wang et al. 2016). For a better identification of BFA bodies, we additionally used the dye FM4-64, which can emit fluorescence in a lipophilic membrane environment. FM4-64 marks the plasma membrane in the first minutes following application to the cell, then may be endocytosed and in the presence of BFA become accumulated in BFA bodies (Bolte et al. 2004). We observed BFA bodies positive for both, HA_3_-TTN5-Alexa 488 and FM4-64 signals (Figure 4B). Similar patterns were observed for YFP-TTN5-derived signals in YFP-TTN5-expressing roots (Figure 4C). Hence, HA_3_-TTN5 and YFP-TTN5 can be present in similar subcellular membrane compartments.

**Figure 4.**
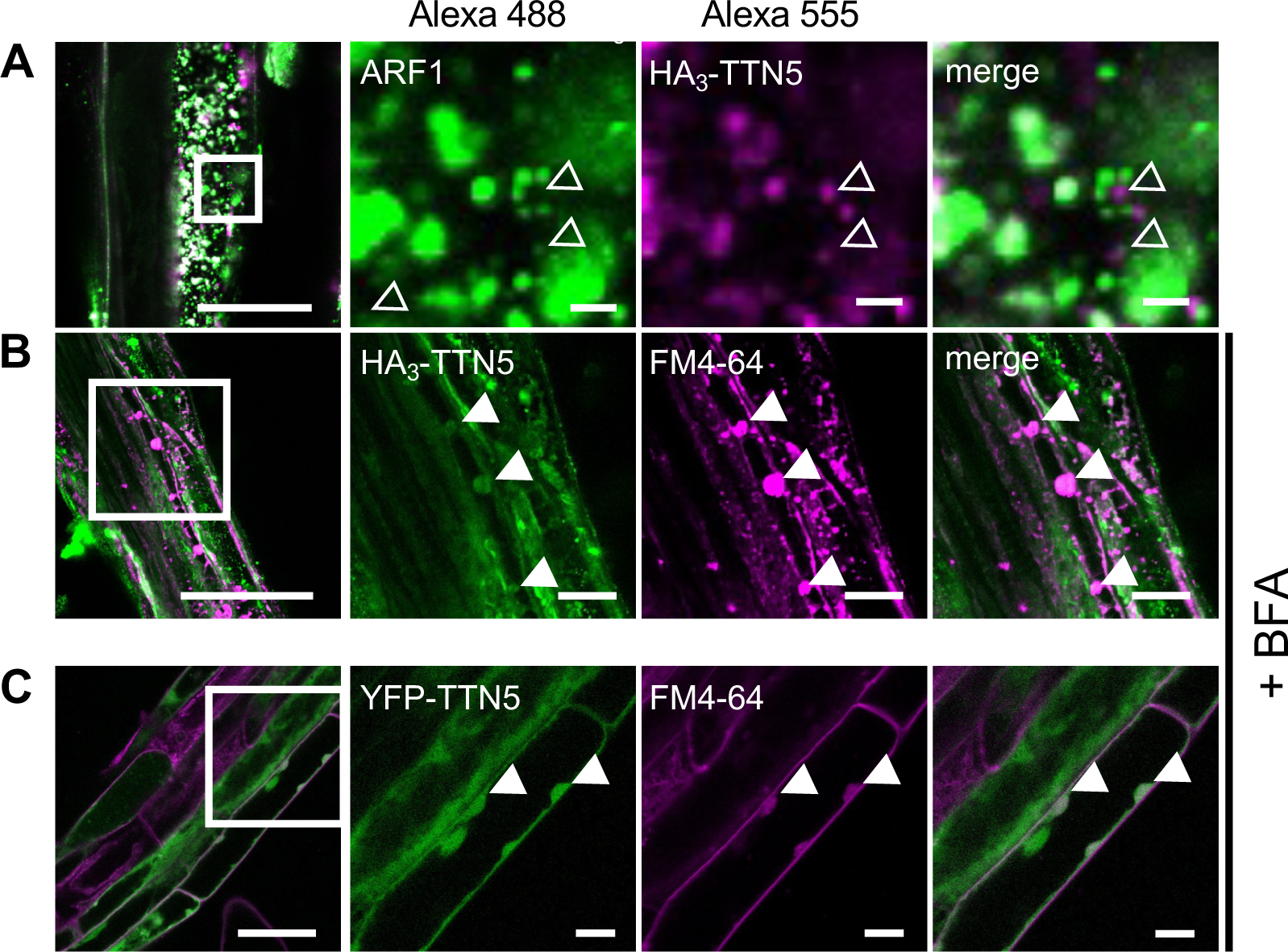
Whole-mount Immunolocalization analysis of HA_3_-TTN5 in Arabidopsis including colocalization with Golgi marker and localization in BFA bodies. (A-B), Representative images showing whole-mount immunostaining of HA_3_-TTN5 seedlings with different types of markers for colocalization analysis. (A), Detection of HA_3_-TTN5 (chicken α-HA primary antibody, Alexa 555-labeled secondary α-chicken antibody) with Golgi and TGN marker ARF1 (rabbit α-ARF1 primary antibody, Alexa-488-labeled secondary α-rabbit antibody). Both fluorescence signals were detected in vesicle-like structures in root cells in close proximity to each other but mostly not colocalizing. The experiment was repeated twice with three seedlings. (B) Detection of HA_3_-TTN5 (rabbit α-HA primary antibody, Alexa-488-labeled secondary α-rabbit antibody) and staining with lipid membrane dye FM4-64 after brefeldin A (BFA) treatment (72 µM, 1 h). Alexa-488 signals colocalized with FM4-64 in BFA bodies in root cells. The experiment was repeated three times with three seedlings. (C), in comparison, YFP fluorescence in YFP-TTN5 seedlings, co-analyzed with FM4-64 after BFA treatment (36 µM, 30 min). YFP fluorescence signals colocalized with FM4-64 in BFA bodies similar as in (B). The experiment was performed once with three independent YFP-TTN5 lines. Colocalizing signals in the two channels are indicated by filled white arrowheads, whereas signals that do not colocalize in the two channels are indicated by empty white arrowheads. Scale bar overview: 50 µm, close-up: 10 µm.

We did not observe any staining in nuclei or ER when performing HA_3_-TTN5 immunostaining (Figure 3P; Figure 4A, B), as was the case for fluorescence signals in YFP-TTN5-expressing cells. Presumably, this can indicate that either the nuclear and ER signals seen with YFP-TTN5 correspond to the smaller proteins detected, as described above, or that immunostaining was not suited to detect them. Hence, we focused interpretation on patterns of localization overlapping between the fluorescence staining with YFP-labeled TTN5 and with HA_3_-TTN5 immunostaining, such as the particular signal patterns in the specific punctate membrane structures.

Taken together, signals of YFP-TTN5 and HA_3_-TTN5 were located in multiple membrane compartments in the epidermis of different Arabidopsis organs and of *N. benthamiana* leaves, including the particular ring-like punctate structures and vesicles. Fluorescence signals in YFP-TTN5 and YFP-TTN5^Q70L^-expressing seedlings displayed high mobility in the cells, as expected from a function of a GTPase in the active state in dynamic processes such as vesicle trafficking. In contrast to that, fluorescence signals in YFP-TTN5^T30N^ were less mobile, in line with the root length phenotype conferred by HA_3_-TTN5 ^T30N^, speaking in favor of the observed kinetics for TTN5^T30N^ with a very fast nucleotide exchange rate and loss of affinity to nucleotides. Altogether, TTN5 intracellular localization seems complex, indicating that TTN5 may have multiple cellular functions as an active GTPase as it can be associated with different intracellular structures of the endomembrane system.

### TTN5 may associate with components of the cellular endomembrane system

The overlapping localization of HA_3_-TTN5 and YFP-TTN5 signals prompted us to better resolve the membrane structures and compartments. The endomembrane system is highly dynamic in the cell. Well-established fluorescent endomembrane markers and pharmacological treatments help to determine the nature of individual components of the system in parallel to colocalization studies with proteins of interest such as TTN5. We conducted the colocalization experiments in *N. benthamiana* leaf epidermis. We just described above that fluorescence signals were comparable between *N. benthamiana* leaf epidermis and Arabidopsis cotyledons or root epidermis. Moreover, it represents an established system for functional association of fluorescent proteins with multiple endomembrane components and optimal identification of membrane structures (Brandizzi et al. 2002, Hanton et al. 2009).

At first, we further investigated the endoplasmic reticulum (ER)-Golgi connection. This site is characteristic of association with small GTPases like the already tested ARF1, involved in COPI vesicle transport from Golgi to the ER (Just and Peränen 2016). This time, we tested another Golgi marker, the soybean (*Glycine max*) protein α-1,2 mannosidase 1 (GmMan1). GmMan1 is a glycosidase that acts on glycoproteins at the *cis*-Golgi, facing the ER (Figure 5A). GmMan1-mCherry-positive Golgi stacks are visible as nearly round punctuate structures throughout the whole cell (Nelson et al. 2007, Wang et al. 2016). Fluorescence signals in leaf discs, transiently expressing YFP-TTN5 and its mutant variants, partially colocalized with GmMan1-mCherry signals at the Golgi stacks (Figure 5B-D). We also observed YFP fluorescence signals in the form of circularly shaped ring structures with a fluorescence-depleted center. These structures can be of vacuolar origin as described for similar fluorescent rings in Tichá et al. (2020) for ANNI-GFP. Further, quantitative analysis reflected the visible colocalization of the GmMan1 marker and YFP fluorescence with Pearson coefficients 0.63 (YFP-TTN5), 0.65 (YFP-TTN5^T30N^) and 0.68 (YFP-TTN5^Q70L^) (Supplementary Figure S8A; see also similar results obtained with overlap coefficients), indicating a strong correlation between the two signals. We performed an additional object-based analysis to compare overlapping YFP fluorescence signals in YFP-TTN5-expressing leaves with GmMan1-mCherry signals (YFP/mCherry ratio) and *vice versa* (mCherry/YFP ratio). We detected 24 % overlapping YFP-fluorescence signals for TTN5 with Golgi stacks, while in YFP-TTN5^T30N^ and YFP-TTN5^Q70L^-expressing leaves, signals only shared 16 and 15 % overlap with GmMan1-mCherry-positive Golgi stacks (Supplementary Figure S8B). Some YFP-signals did not colocalize with the GmMan1 marker. This effect appeared more prominent in leaves expressing YFP-TTN5^T30N^ and less for YFP-TTN5^Q70L^, compared to YFP-TTN5 (Figure 5B-D). Indeed, we identified 48 % GmMan1-mCherry signal overlapping with YFP-positive structures in YFP-TTN5^Q70L^ leaves, whereas 43 and only 31 % were present with YFP fluorescence signals in YFP-TTN5 and YFP-TTN5^T30N^-expressing leaves, respectively (Supplementary Figure S8B), indicating a smaller amount of GmMan1-positive Golgi stacks colocalizing with YFP signals for YFP-TTN5^T30N^. Hence, the GTPase-active TTN5 forms are likely more present at *cis*-Golgi stacks compared to TTN5^T30N^.

**Figure 5:**
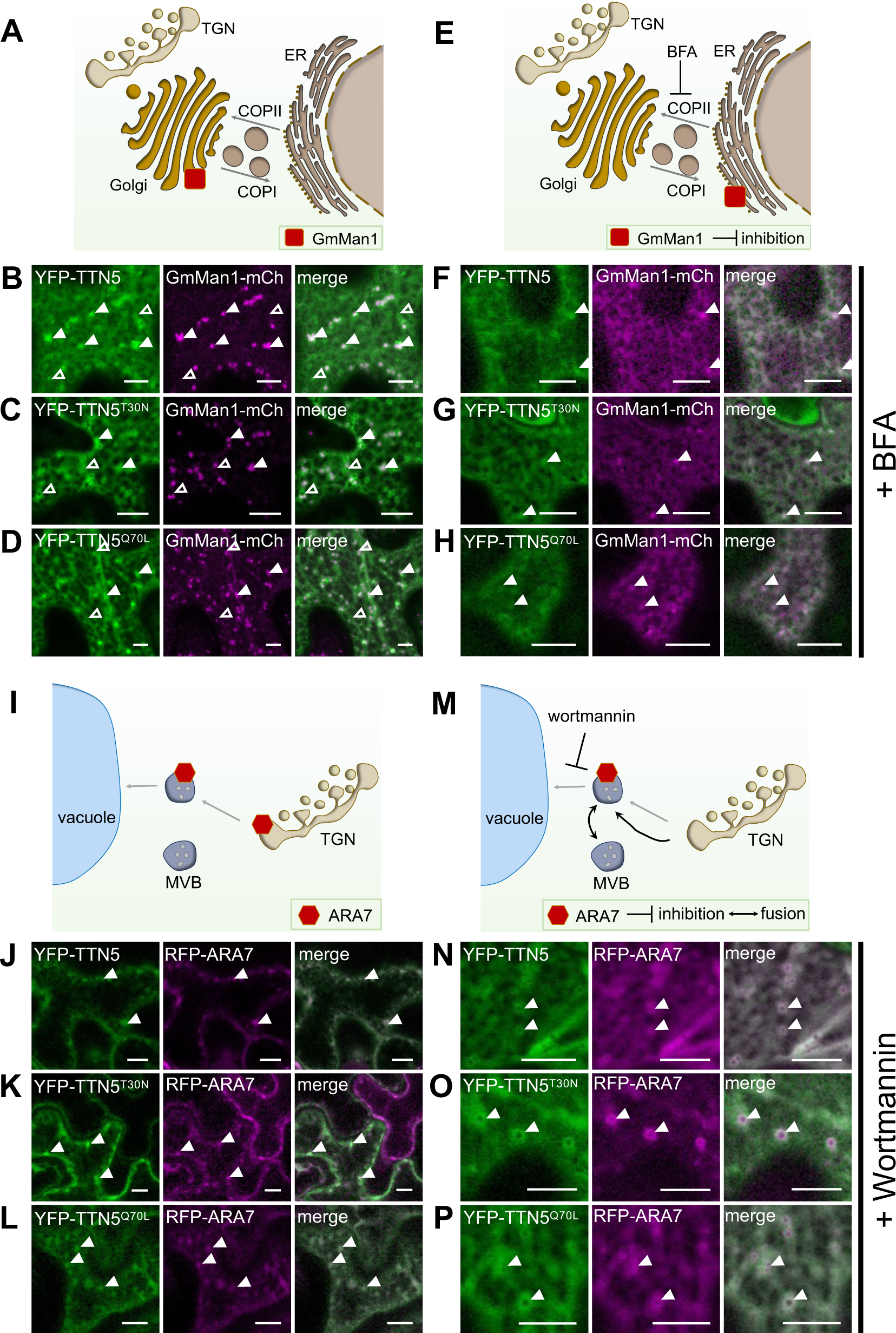
TTN5 may be associated with the endomembrane system in *N. benthamiana* leaf epidermal cells. YFP fluorescence signals were localized in *N. benthamiana* leaf epidermal cells transiently transformed to express YFP-TTN5, YFP-TTN5^T30N^ and YFP-TTN5^Q70L^ *via* fluorescent confocal microscopy. Specific markers indicating the endomembrane system were used. (A), Schematic representation of GmMan1 localization at the *cis*-Golgi site. (B-D), Partial colocalization of YFP signal with the Golgi marker GmMan1-mCherry at *cis*-Golgi stacks (filled white arrowheads). Additionally, YFP fluorescent signals were detected in non-colocalizing punctate structures with depleted fluorescence in the center (empty white arrowheads). (E), Schematic representation of GmMan1 localization at the ER upon brefeldin A (BFA) treatment. BFA blocks ARF-GEF proteins which leads to a loss of Golgi *cis*-cisternae and the formation of BFA-induced compartments due to an accumulation of Golgi stacks up to a redistribution of the Golgi to the ER by fusion of the Golgi with the ER (Renna and Brandizzi 2020). (F-H), Redistribution of Golgi stacks was induced by BFA treatment (36 µM, 30 min). GmMan1-mCherry and YFP fluorescence signals were present in the ER and in colocalizing punctate structures. (I), Schematic representation of ARA7 localization at the *trans*-Golgi network (TGN) and multi-vesicular bodies (MVBs). (J-L), Colocalization of YFP fluorescence signal with the MVB marker RFP-ARA7. (M), Schematic representation of ARA7 localization in swollen MVBs upon wortmannin treatment. Wortmannin inhibits phosphatidylinositol-3-kinase (PI3K) function leading to the fusion of TGN/EE to swollen MVBs (Renna and Brandizzi 2020). (N-P), MVB swelling was obtained by wortmannin treatment (10 µM, 30 min). ARA7-RFP was colocalizing with YFP signal in these swollen MVBs. Chemical treatment-induced changes were imaged after 25 min incubation. Colocalization is indicated with filled arrowheads, YFP signal only with empty ones. Corresponding colocalization analysis data is presented in Supplementary Figure S8. Scale bar 10 μm.

Next, we evaluated the Golgi localization by BFA treatment. The action of BFA causes a corresponding redistribution of GmMan1-mCherry (Ritzenthaler et al. 2002, Wang et al. 2016) (Figure 5E). We found that upon BFA treatment, GmMan1-mCherry signal was present in the ER and in BFA-induced compartments. YFP-signal of YFP-TTN5 constructs showed partially matching localization with GmMan1-mCherry upon BFA treatment suggesting a connection of TTN5 to Golgi localization (Figure 5F-H). Hence, the colocalization with GmMan1-mCherry and BFA treatment was indicative of YFP signals localizing to Golgi stacks upon YFP-TTN5 expression, while the lower association of mostly the YFP-TTN5^T30N^ mutant form with this membrane compartment was noted.

Second, we investigated localization to the endocytic compartments, endosomes of the *trans*-Golgi network (TGN) and multivesicular bodies (MVBs) using the marker RFP-ARA7 (RABF2B), a small RAB-GTPase present there (Kotzer et al. 2004, Lee et al. 2004, Stierhof and El Kasmi 2010, Ito et al. 2016) (Figure 5I). These compartments play a role in sorting proteins between the endocytic and secretory pathways, with MVBs developing from the TGN and representing the final stage in transport to the vacuole (Valencia et al. 2016, Heucken and Ivanov 2018). Colocalization studies revealed that YFP signal in YFP-TTN5-expressing leaves was present at RFP-ARA7-positive MVBs (Figure 5J). Noticeably, overlaps between RFP-ARA7 and YFP fluorescence signals upon TTN5^T30N^ expression were lower than for the other TTN5 forms (Figure 5J-L; Supplementary Figure S8C, D). We obtained a Pearson coefficient for the pair of either YFP fluorescence upon YFP-TTN5 or YFP-TTN5^Q70L^ expression with RFP-ARA7 of 0.78, whereas a coefficient of only 0.59 was obtained with YFP-TTN5^T30N^ confirming the visual observation (Supplementary Figure S8C; see also similar results for overlap coefficients). Object-based analysis showed that, RFP-ARA7-positive structures had an overlap with YFP fluorescence in YFP-TTN5-expressing (29 %) leaves and even more with YFP-TTN5^Q70L^ (75 %) signals unlike with YFP-TTN5^T30N^ signals (21 %) (Supplementary Figure S8D). Based on this, signals of YFP-TTN5^Q70L^ and YFP-TTN5 tended to colocalize better with ARA7-positive compartments than YFP-TTN5^T30N^.

To test MVB localization, we treated plant cells with wortmannin, a common approach to studying endocytosis events. Wortmannin is a fungal metabolite that inhibits phosphatidylinositol-3-kinase (PI3K) function and thereby causes swelling of the MVBs (Cui et al. 2016) (Figure 5M). RFP-ARA7-expressing cells showed the expected typical wortmannin-induced formation of doughnut-like shaped MVBs (Jaillais et al. 2008). The YFP fluorescence signals in YFP-TTN5-expressing leaves partially colocalized with these structures (Figure 5N-P) indicating that fluorescence signals upon YFP-TTN5 expression and the two mutants are present in MVBs. YFP signals in YFP-TTN5Q70L-expressing leaf discs were located even to a greater extent to MVBs than in wild-type YFP-TTN5 and much more than in YFP-TTN5^T30N^-expressing cells, suggesting an active role of YFP-TTN5^Q70L^ in MVBs, for example in the lytic degradation pathway or the recycling of proteins, similar to ARA7 (Kotzer et al. 2004).

Finally, to investigate a possible connection of TTN5 with the plasma membrane, we colocalized YFP signals of the YFP-TTN5 constructs with the dye FM4-64, which can emit fluorescence in a lipophilic membrane environment and marks the plasma membrane in the first minutes following application to the cell (Bolte et al. 2004) (Figure 6A). Fluorescence signals for all three forms of TTN5 colocalized with FM4-64 at the plasma membrane in a similar manner (Figure 6B-D). To further investigate plasma membrane localization, we performed mannitol-induced plasmolysis. YFP signals for all three YFP-TTN5 constructs were then located similarly to FM4-64-stained Hechtian strands, thread-like structures attached to the apoplast visible upon plasmolysis and surrounded by plasma membrane (Figure 6E-G).

**Figure 6.**
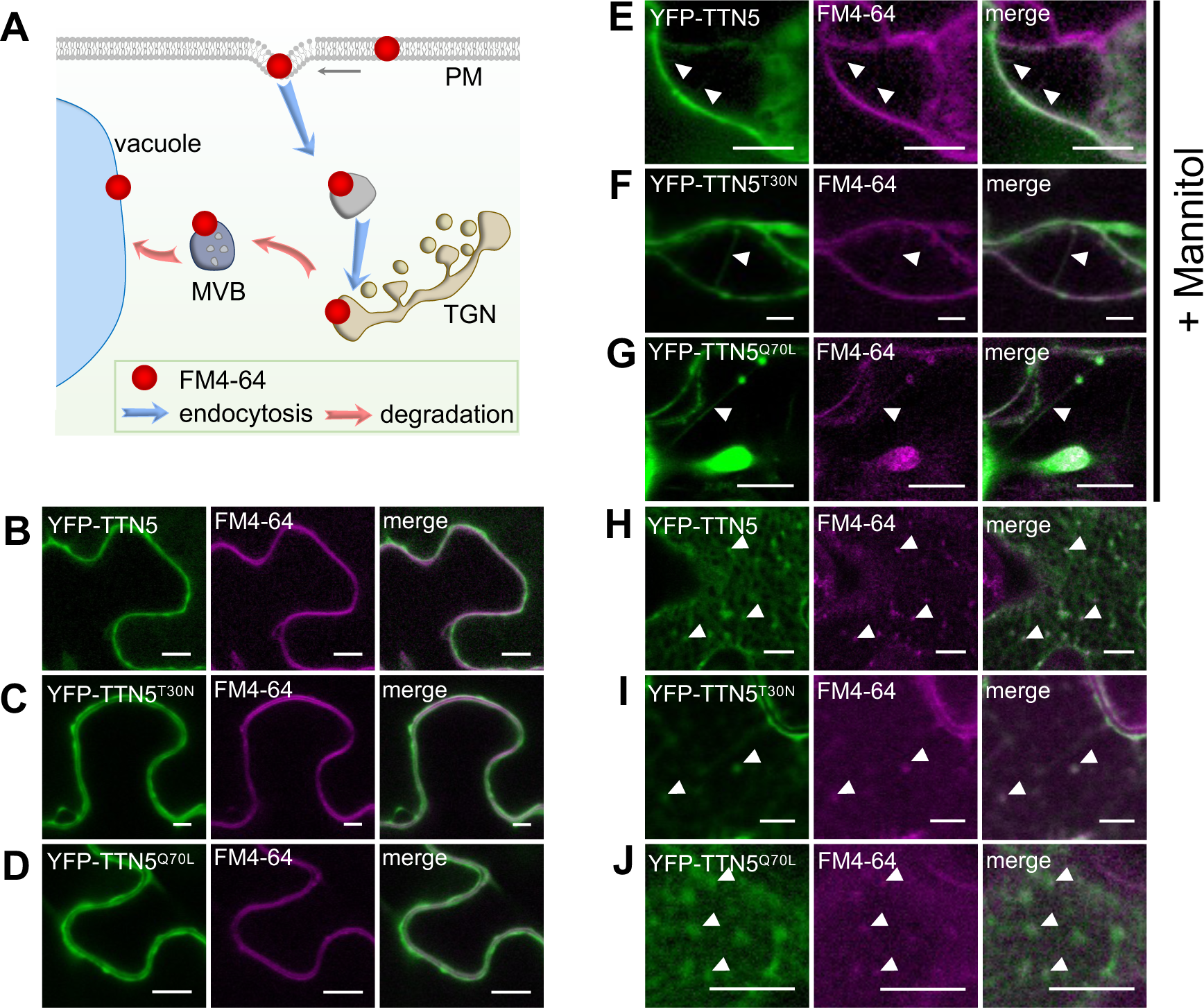
TTN5 may colocalize with endocytosed plasma membrane material. (A), Schematic representation of progressive stages of FM4-64 localization and internalization in a cell. FM4-64 is a lipophilic substance. After infiltration, it first localizes in the plasma membrane, at later stages it localizes to intracellular vesicles and membrane compartments. This localization pattern reflects the endocytosis process (Bolte et al. 2004). (B-J), YFP fluorescence signals were localized in *N. benthamiana* leaf epidermal cells together with the plasma membrane dye FM4-64 *via* fluorescent confocal microscopy, following transient transformation to express YFP-tagged TTN5, YFP-TTN5^T30N^ and YFP-TTN5^Q70L^. (B-D), YFP signals colocalized with FM4-64 at the plasma membrane. (E-G), Plasma membrane localization of YFP fluorescence was evaluated after mannitol-induced (1 M) plasmolysis. The formation of Hechtian strands is a sign of plasma membrane material and fluorescence staining there is indicated with filled arrowheads. (H-J), Internalized FM4-64 was present in vesicle-like structures that showed YFP signals. Colocalization is indicated with filled arrowheads. Scale bar 10 μm.

**Figure 7:**
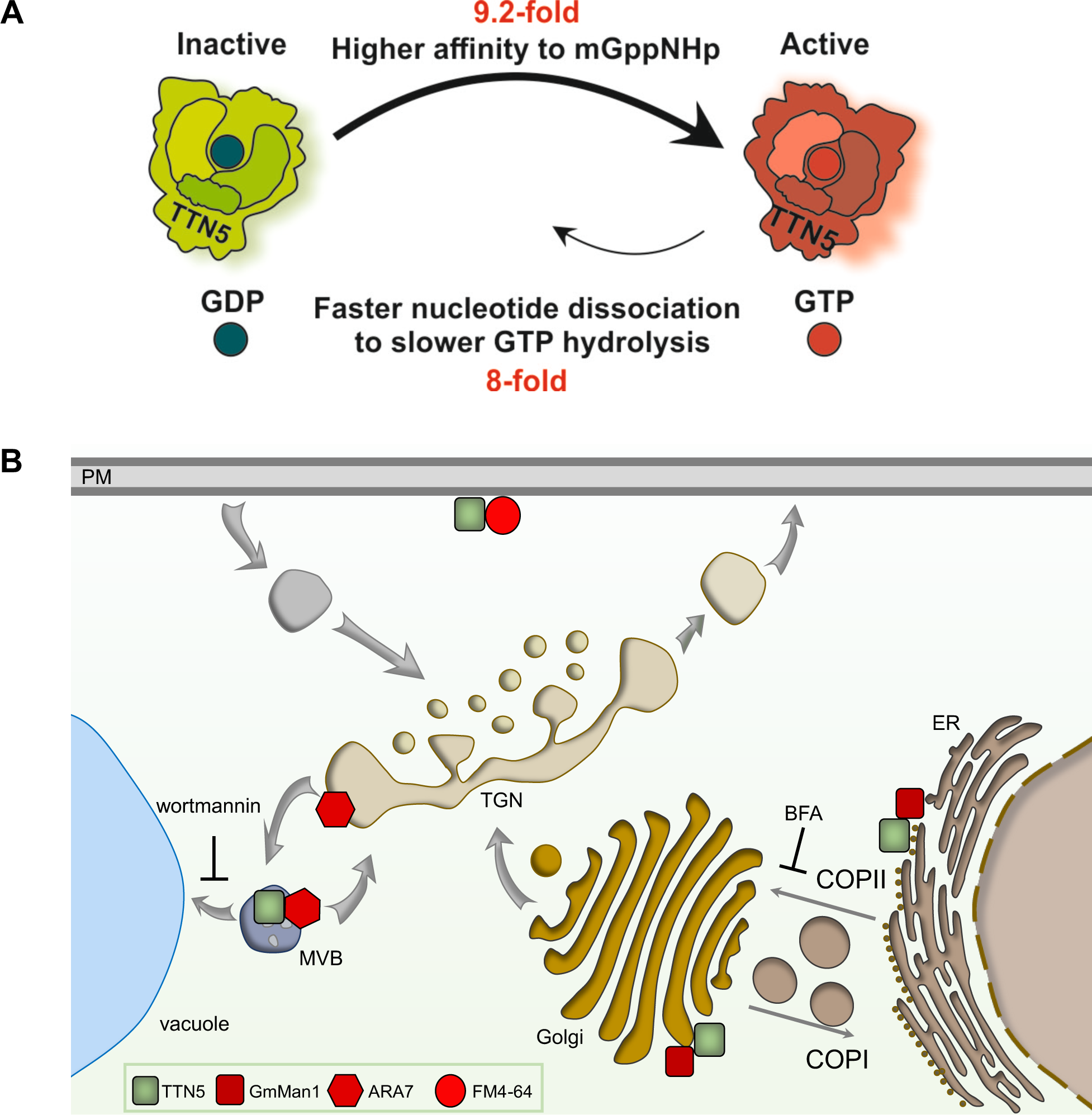
Schematic models summarize TTN5 kinetic GTPase activities and potential localization within the cell. (A), Model of the predicted GTPase nucleotide exchange and hydrolysis cycle mechanism of TTN5 based on the biochemical investigation. TTN5 affinity to mGppNHp was 9.2-fold higher compared to mGDP resulting in a fast switching from an inactive GDP-loaded form to an active GTP-loaded one. mGppNHp dissociation was 8-fold faster as GTP hydrolysis but both processes were much slower than nucleotide association. TTN5 kinetics identified it as a non-classical GTPase which tended to stay in a GTP-loaded form even under resting conditions. (B), Presumed TTN5 locations within the cell. TTN5 (green square) can be present at the plasma membrane (PM) similar as FM4-64 (red circle) or in the endomembrane compartments of the *trans*-Golgi network (TGN) or multivesicular body (MVB) as found by colocalization with ARA7 (red hexagon). Additionally TTN5 might colocalize with GmMan1-positive (red square) Golgi stacks.

In summary, these colocalization experiments showed that YFP signals upon YFP-TTN5 expression were found in different membrane sites of the endomembrane system, including Golgi, MVBs and plasma membrane. We figured that similar to other ARF proteins, this pattern can indicate that TTN5 might participate in a highly dynamic vesicle trafficking process. Indeed, when we recorded the dynamic movement of YFP signals inside *N. benthamiana* leaf epidermis cells, YFP-TTN5 and YFP-TTN5^Q70L^ derived signals colocalized with GmMan1-mCherry and revealed high motion over time, while, again, this was less the case for the YFP-TTN5^T30N^ construct (Supplementary Video Material S1M-O).

One potential cellular trafficking route is the degradation pathway to the vacuole. We, therefore, investigated fluorescence localization upon YFP-TTN5 transient expression in late endosomal compartments that might be involved in vacuolar targeting. FM4-64 is used as a marker for membranes of late endosomal compartment and vacuole targeting, since following plasma membrane visualization FM4-64-stained endocytic vesicles become apparent at later stages as well as vacuolar membrane staining (Ueda et al. 2001, Emans et al. 2002, Dhonukshe et al. 2007, Ivanov and Vert 2021). Hence, we colocalized YFP signals with FM4-64-positive compartments at later time points. Next to colocalization of YFP fluorescence in YFP-TTN5-expressing leaves with FM4-64 at the plasma membrane, we detected colocalization with fluorescent compartments in the cell, which was similar for the two mutant forms (Figure 6H-J). This indicates that YFP-TTN5 may be involved in the targeting of endocytosed plasma membrane material, irrespective of the mutations.

In summary, YFP signals upon YFP-TTN5 and YFP-TTN5^Q70L^ expression were dynamic and colocalized with endomembrane structures, whereas fluorescence signal in YFP-TTN5^T30N^-expressing leave discs tended to be less mobile and dynamic and colocalized less.

## Discussion

This work provides evidence that the small ARF-like GTPase TTN5 has a very rapid intrinsic nucleotide exchange capacity with a conserved nucleotide switching mechanism. TTN5 might be primarily present in a GTP-loaded active form in a cell. TTN5 might be also a dynamic protein inside cells with respect to its localization to membrane structures, which can be a hint on association with vesicle transport and different processes of the endomembrane system. The active TTN5^Q70L^ mutant was capable of nucleotide switching and might be mostly similarly localized as TTN5 in a cell. The TTN5^T30N^ mutant, on the other hand, was affected by a lower nucleotide exchange capacity than the other TTN5 forms. It differed significantly in localization properties and its dynamics, albeit depending on cell types. TTN5^T30N^ also conferred a root length phenotype. Therefore, the GTP-bound state that we presume for TTN5 is most likely very critical for protein localization and dynamics in cells.

### TTN5 exhibits characteristic GTPase functions

TTN5 was classified as an ARL2 homolog of the ARF GTPases based on its sequence similarity. The sequence analysis suggested nucleotide binding (McElver et al. 2000) which is reinforced by structural prediction suggesting the formation of a nucleotide-binding pocket by the binding motifs. Nucleotide association and dissociation of TTN5, TTN5^T30N^ and TTN5^Q70L^ indicated that TTN5 along with the two mutant forms can bind guanine nucleotides. The k_on_ values of TTN5^T30N^ and TTN5 were nearly the same, indicating no effect of the mutation on the GDP-binding characteristics as was expected in the absence of a GEF. The k_on_ value for TTN5^Q70L^ was clearly higher than that of the wild-type form, indicating that this mutant can bind GDP faster than TTN5 to form the nucleotide-bound form. Compared with other members of the Ras superfamily, it was in the range of HRAS (Hanzal-Bayer et al. 2005) and around ten times slower than the fast association of RAC1 (Jaiswal et al. 2013). Intrinsic nucleotide exchange measurements of TTN5 and TTN5^Q70L^ have shown remarkably fast nucleotide exchange rates, when compared to other well-studied RAS proteins. The intrinsic nucleotide exchange reaction rates for RAC1, RAC2 and RAC3 have been mentioned around 40.000 s (Haeusler et al. 2006). Our data show that TTN5 is faster in nucleotide exchange rate and very similar to that of human ARL2 (Hanzal-Bayer et al. 2005, Veltel et al. 2008). This suggests that TTN5 quickly replaces GDP for GTP and transforms from an inactive to an active state. This behavior indicates that TTN5 presumably should not require interaction with GEFs for activation in cells. This can also be an explanation for the case of TTN5^Q70L^. Small GTPases with substitutions of the glutamine of the switch II region (*e.g*., Glu-71 for HsARF1 and ARL1, Glu-61 for HRAS) are constitutively active (Zhang et al. 1994, Van Valkenburgh et al. 2001, Karnoub and Weinberg 2008). Accordingly, TTN5^Q70L^ is likely to exchange GDP rapidly to GTP and switch itself to stay in an active form as suggested by the fast intrinsic nucleotide exchange rate. Interestingly, TTN5^T30N^ resulted in an even higher dissociation rate constant k_off_. The calculated K_d_ confirmed the higher nucleotide-binding affinity for GDP of TTN5 and TTN5^Q70L^ compared with TTN5^T30N^. Reports on human ARL2, ARF6 and ARL4D showed that their corresponding T30N mutants led to a decreased affinity to GDP similar to TTN5^T30N^ (Macia et al. 2004, Hanzal-Bayer et al. 2005, Li et al. 2012).

Interestingly, a comparison of mdGDP with mGppNHp revealed a higher GTP affinity for all three versions, with the highest for TTN5^Q70L^. These high GTP affinities in combination with the fast GDP exchange rates and extremely slow hydrolysis pinpointed to a GTP-loaded TTN5 even in the rested state, which is very uncommon for small GTPases. This atypical behavior is already reported for a few non-classical RHO GTPases like RHOD or RIF (Jaiswal et al. 2013). This unusual GTP-bound active state along with the lacking *N*-myristoylation and phylogenetic distances (Boisson et al. 2003, Vernoud et al. 2003) strengthens that there are major differences between TTN5 and other ARF family members. The similarity between the wild type and TTN5^Q70L^ is consistent with the previous report on human ARL2 in which wild-type and Q70L proteins showed only a little difference in binding affinity (Hanzal-Bayer et al. 2005). Additionally, an equivalent ratio of nucleotide affinity was found between HRAS and HRAS^Q61L^, but with a much higher affinity typical for small GTPases (Der et al. 1986). Since Gln-70 at the switch II region is important for GAP-stimulated GTP hydrolysis (Cherfils and Zeghouf 2013), we assume that nucleotide exchange activity is unaffected by this amino acid substitution.

To date, no GEF protein for TTN5 is reported. The Arabidopsis genome encodes only two of the five mammalian GEF subgroups, namely the large ARF-GEF subgroups, the BFA-inhibited GEF (BIG) and the Golgi Brefeldin A (BFA)-resistance factor 1 (GBF/GNOM) family (Memon 2004, Wright et al. 2014, Brandizzi 2018). Potential interactions with these proteins are of high interest and can also point to functions of TTN5 as a co-GEF as it is proposed for HsARL3 and HsARL2 with their effector BART by stabilizing the active GTPase (ElMaghloob et al. 2021). Especially, interactions at the nucleotide-binding site, which are prevented in the TTN5^T30N^ mutant, will be of great interest to study further functions and interaction partners of TTN5.

Taken together, the categorization as a non-classical GTPase has three implications: First, the very slow hydrolysis rate predicts the existence of a TTN5-GAP. Second, TTN5^T30N^ may function as a dominant-negative mutant and in the presence of a GEF, it cannot bind GDP. Third, the TTN5^Q70L^ hydrolysis rate is not decreased.

### TTN5 may act in the endomembrane system

The localization data on YFP- and HA_3_-TTN5 suggest that it may be localized at different cellular membrane compartments which is typical for the ARF-like GTPase family (Memon 2004, Sztul et al. 2019) and supports potential involvement of TTN5 in endomembrane trafficking. We based the TTN5 localization data on tagging approaches with two different detection methods to enhance reliability of specific protein detection. Even though YFP-TTN5 did not complement the embryo-lethality of a *ttn5* loss of function mutant, we made several observations that suggest YFP-TTN5 signals to be meaningful at various membrane sites. YFP-TTN5 may not complement due to differences in TTN5 levels and interactions in some cell types, which were hindering specifically YFP-TTN5 but not HA_3_-TTN5. In a previous study, overexpression of ARF1 did not affect intracellular localization compared to endogenous tagged-ARF1 but differed in function to form tubulated structures (Bottanelli et al. 2017). Though constitutively driven, the YFP-TTN5 expression may be delayed or insufficient at the early embryonic stages resulting in the lack of embryo-lethal complementation. On the other hand, the very fast nucleotide exchange activity may be hindered by the presence of a large YFP-tag in comparison with the small HA_3_-tag which is able to rescue the embryo-lethality. The lack of complementation represents a challenge for the localization of small GTPases with rapid nucleotide exchange in plants. Despite of these limitations, we made relevant observations in our data that made us believe that YFP signals in YFP-TTN5-expressing cells at membrane sites can be meaningful. At first, using pharmacological treatments and colocalization with known membrane compartment markers, we noted that various particular membrane compartments showed YFP signals, such as the punctate small ring-like structures resembling previously reported ANNI-GFP staining (Tichá et al. 2020), large ring-like structures resulting from wortmannin treatment and BFA bodies, all of which are meaningful for studies of vesicle transport and plasma membrane protein regulation processes (Wang et al. 2009, Suo et al. 2021). Furthermore, the fluorescence signals obtained with YFP-TTN5 constructs also depended on T30 and Q70 residues. Point mutant YFP-TTN5 forms, and particularly the YFP-TTN5^T30N^ had partly quite distinct fluorescence localization patterns, such as reduced mobility in certain cells and differing degrees of colocalization with the utilized markers. Next to this, HA_3_-TTN5^T30N^ seedlings showed reduced root growth which may be due to the same reasons as the altered localization and mobility. Since TTN5^T30N^ has, based on the enzyme kinetic results, a very fast nucleotide exchange rate and lost affinity to nucleotides compared to TTN5, these differing YFP fluorescence patterns of the YFP-TTN5^T30N^ construct at membrane sites and the effect on root growth are not unexpected to occur. Hence, we considered these specific YFP localization signals at membrane sites for valid interpretation, especially when supported by HA_3_-TTN5 immunodetection.

Following up, colocalization analysis showed that both *cis*-Golgi and MVB-positive structures colocalized to a higher proportion with YFP signals of the YFP-TTN5^Q70L^ construct compared with signals in YFP-TTN5T30N-expressing cells. This could be an indicator of the site of TTN5 action, considering our knowledge of the activation of ARF GTPases and ARL proteins in other organisms which show high TTN5 sequence similarity. They are usually recruited or move to their place of action upon interacting with their specific GEF, which leads to GDP to GTP exchange-dependent activation (Sztul et al. 2019, Nielsen 2020, Adarska et al. 2021). Though our biochemical data implies no need for a typical GTPase-GEF interaction for activation, GEF interaction can be still important for the localization. Most of the effector-GTPase interactions take place in their GTP-bound form (Sharer and Kahn 1999, Hanzal-Bayer et al. 2005). One exception is the role of TTN5 sequence-based homologs in microtubule dynamics. ARL2/Alp41-GDP interacts with Cofactor D/Alp1^D^ (Bhamidipati et al. 2000, Mori and Toda 2013). Another possibility is a hindrance of dimerization by the T30N mutation. ARF1 protein dimer formation is important for the formation of free vesicles (Beck et al. 2009, Beck et al. 2011) associated with cell mobility which was disturbed in YFP-TTN5^T30N^-expressing cells. The colocalization of YFP fluorescence in YFP-TTN5-expressing cells with ARA7-positive structures even still in the wortmannin-induced swollen state, triggered by the homotypic fusion of MVBs (Wang et al. 2009), may indicate that TTN5 performs similar functions in relation to ARA7. ARA7 is involved in cargo transport in the endocytic pathway to the vacuole, with a role, for example, in the endocytosis of plasma membrane material (Ueda et al. 2001, Sohn et al. 2003, Kotzer et al. 2004, Ebine et al. 2011). The colocalization of FM4-64-labeled endocytosed vesicles with fluorescence in YFP-TTN5-expressing cells may indicate that TTN5 is involved in endocytosis and the possible degradation pathway into the vacuole. Our data on colocalization with the different markers support the hypothesis that TTN5 may have functions in vesicle trafficking.

A potential explanation of the YFP localization to similar compartments in YFP-TTN5- and YFP-TTN5^Q70L^-expressing cells compared to fluorescence signal of YFP-TTN5^T30N^ expression can be based on a special feature of TTN5 in the ARF family. ARF GTPases are mostly myristoylated on Gly-2, which is essential for their membrane binding. TTN5 as well as ARL2 and ARL3 lack this myristoylation though Gly-2 is present (Boisson et al. 2003, Kahn et al. 2006). ARL2 and ARL3 are still able to bind membranes, probably only by their N-terminal amphipathic helix as it was established for SAR1, with an ARL2 membrane-binding efficiency being nucleotide-independent (Lee et al. 2005, Kapoor et al. 2015). We suggest similar behavior for TTN5, as detected YFP signals localized to membranous compartments. Based on the varying colocalization degrees, with the fluorescence signals of YFP-TTN5^T30N^ construct being less prominent at the Golgi and MVBs, compared to YFP-TTN5 and YFP-TTN5^Q70L^, we hypothesize that different membrane localization could be associated with a nucleotide- or nucleotide exchange-dependent process. In a nucleotide-free or GDP-bound state, TTN5 may be predominantly present close to the plasma membrane, while in an active GTP-bound state, which according to enzyme kinetics should be the regular one, is dynamically linked with the endomembrane system. Interestingly, with respect to the intracellular dynamics, we observed that the TTN5^T30N^ mutant had a different behavior in different organ types. This could be due to differing GEFs being differentially expressed. Likewise, it is conceivable that the constitutively expressed TTN5 has different effector binding partners.

This broad diversity of biological functions of proteins with high sequence similarity to TTN5 associated with a variety of signaling cascades is also reflected by very different protein partners for that. Few orthologs of human ARL2 interaction partners are present in Arabidopsis. It is therefore exceedingly interesting to identify interacting proteins to determine whether TTN5 performs similar functions as HsARL2 or what other role it may play. Such interactions might also explain why TTN5 is essential in plants with regard to a potential GTP-dependence for TTN5 function which fits to already known functions of other ARF GTPases (Sztul et al. 2019, Nielsen 2020, Adarska et al. 2021). In addition, ARF proteins are affected by a similar set of GEFs and GAPs, indicating an interconnected network in ARF signaling. ARF double knockdowns revealed specific phenotypes, suggesting redundancy in the ARF family (Volpicelli-Daley et al. 2005, Kondo et al. 2012, Nakai et al. 2013, Adarska et al. 2021). The investigation of the TTN5 connection in the ARF family might reveal a missing link in ARF signaling and cell traffic.

## Conclusion

In this study, we identified TTN5 as a functional GTPase of the ARF-like family. TTN5 had not only sequence similarity with human ARL2 but also, both these two proteins share a very rapid nucleotide exchange capacity in contrast to other characterized ARF/ARL proteins. TTN5 has a faster nucleotide dissociation rate to a slower GTP hydrolysis rate and a higher affinity to GTP compared to GDP. Thus, TTN5 is a non-classical GTPase that most likely accumulates in a GTP-bound state in cells in line with certain cellular phenotypes and protein localization data. The nucleotide exchange capacity affected the localization and dynamics of YFP-tagged TTN5 protein forms and associated TTN5 with the endomembrane system. In the future, the identification of potential TTN5 GEF, GAP and effector proteins as well as other interaction partners and particularly potential plasma membrane target proteins as cargo for vesicle transport will be of great interest to clarify the potential roles of TTN5 in endomembrane trafficking and whole-plant physiological contexts.

## Material & Methods

### Arabidopsis plant material and growth conditions

The Arabidopsis *ttn5-1* mutant was previously described (McElver et al. 2000). Heterozygous seedlings were selected by genotyping using the primers TTN5 intron1 fwd and pDAP101 LB1 (Supplementary Table S1). For pro35S::YFP-TTN5 and pro35S::HA_3_-TTN5 constructs, *TTN5*, *TTN5^T30N^*and *TTN5^Q70L^* coding sequences were amplified with B1 and B2 attachment sites for Gateway cloning (Life Technologies) using the primer TITAN5 n-ter B1 and TITAN5 stop B2 (Supplementary Table S1). The obtained PCR fragments were cloned *via* BP reaction (Life Technologies) into pDONR207 (Invitrogen). pro35S::YFP-TTN5 and pro35S::HA_3_-TTN5 constructs were created *via* LR reaction (Life Technologies) with the destination vector pH7WGY2 (Karimi et al. 2005) and pALLIGATOR2 (Bensmihen et al. 2004), respectively. Agrobacteria were transformed with obtained constructs and used for stable Arabidopsis transformation (adapted by (Clough and Bent 1998). Arabidopsis seeds were sterilized with sodium hypochlorite solution (6 % Sodium hypochlorite and 0.1 % Triton X-100) and stored for 24 hours at 4°C for stratification. Seedlings were grown upright on half-strength Hoagland agar medium (1.5 mM Ca(NO_3_)_2_, 0.5 mM KH_2_PO_4_, 1.25 mM KNO_3_, 0.75 mM MgSO_4_, 1.5 µM CuSO_4_, 50 µM H_3_BO_3_, 50 µM KCl, 10 µM MnSO_4_, 0.075 µM (NH_4_)_6_Mo_7_O_24_, 2 µM ZnSO_4_, 50 μM FeNaEDTA and 1 % sucrose, pH 5.8, supplemented with 1.4 % Plant agar (Duchefa)] in growth chambers (CLF Plant Climatics) under long-day condition (16 hours light at 21°C, 8 hours darkness at 19°C). Seedlings were grown for six days (six-day system) or 10 days (10-day system) or 17 days with the last three days on fresh plates (two-week system).

Root length measurement were performed using JMicroVision: Image analysis toolbox for measuring and quantifying components of high-definition images. Version 1.3.4 (https://jmicrovision.github.io, Roduit, N.)

*Nicotiana benthamiana* plants were grown on soil for 2-4 weeks in a greenhouse facility under long-day conditions (16 hours of light, 8 hours of darkness).

### Point mutant generation of *TTN5*

pDONR207:TTN5 was used as a template for site-directed *TTN5* mutagenesis. Primers T5T30Nf and T5T30Nr (Supplementary Table S1) were used to amplify the entire vector generating the TTN5^T30N^ coding sequence and primers TQ70Lf and T5Q70Lr (Supplementary Table S1) were used to amplify the entire vector generating the TTN5^Q70L^ coding sequence. The PCR amplifications were run using the following conditions: 95°C, 30 s; 18 cycles of 95°C, 30 s/ 55°C, 1 min/ 72°C 8 min; 72°C, 7 min. The completed reaction was treated with 10 units of DpnI endonuclease for 1 h at 37°C and then used for *E. coli* transformation. Successful mutagenesis was confirmed by Sanger sequencing.

### *In vitro* GTPase activity assays

An overview of protein expression and purification is shown in Supplementary Figure S2A. Recombinant pGEX-4T-1 bacterial protein expression vectors (Amersham, Germany) containing coding sequences for *TTN5*, *TTN5^T30N^*and *TTN5^Q70L^* were transferred into *E. coli* BL21 (DE3) Rosetta strain (Invitrogen, Germany). Following induction of GST-TTN5 fusion protein expression according to standard procedures. Cell lysates were obtained after cell disruption with a probe sonicator (Bandelin sonoplus ultrasonic homogenizer, Germany) using a standard buffer (300 mM NaCl, 3 mM Dithiothreitol (DTT), 10 mM MgCl_2_, 0.1 mM GDP, 1 % Glycerol and 50 mM Tris-HCl, pH 7.4). GST-fusion proteins were purified by loading total bacterial lysate on a preequilibrated glutathione Sepharose column (Sigma, Germany) using fast performance liquid chromatography system (Cytiva, Germany) (Step 1, affinity-purified GST-TTN5 protein fraction). GST-tagged protein fractions were incubated with thrombin (Sigma, Germany) at 4°C overnight for cleavage of the GST-tag (Step 2, GST cleavage) and applied again to the affinity column (Step 3, yielding TTN5 protein fraction). Purified proteins were concentrated using 10 kDa ultra-centrifugal filter Amicon (Merck Millipore, Germany). The quality and quantity of proteins were analyzed by SDS-protein gel electrophoresis (Bio-Rad), UV/Vis spectrometer (Eppendorf, Germany) and high-performance liquid chromatography (HPLC) using a reversed-phase C18 column (Sigma, Germany) and a pre-column (Nucleosil 100 C18, Bischoff Chromatography) as described (Eberth and Ahmadian 2009) (Supplementary Figure S2B-D).

Nucleotide-free TTN5 protein was prepared from the TTN5 protein fraction (Eberth and Ahmadian 2009) as illustrated in Supplementary Figure S2E. 0.5 mg TTN5 protein was combined with 1 U of agarose bead-coupled alkaline phosphatase (Sigma Aldrich, Germany) for degradation of bound GDP to GMP and Pi in the presence of 1.5-fold molar excess of non-hydrolyzable GTP analog GppCp (Jena Bioscience, Germany). After confirmation of GDP degradation by HPLC, 0.002 U snake venom phosphodiesterase (Sigma Aldrich, Germany) per mg TTN5 was added to cleave GppCp to GMP, G and Pi. The reaction progress of degradation of nucleotides was analyzed by HPLC using 30 µM TTN5 in 30 µl injection volume (Beckman Gold HPLC, Beckman Coulter). After completion of the reaction, in order to remove the agarose bead-coupled alkaline phosphatase, the solution was centrifuged for 10 min at 10000 *g*, 4°C, which was followed by snap freezing and thawing cycles to inactivate the phosphodiesterase. mdGDP (2-deoxy-3-O-N-methylanthraniloyl GDP)- and mGppNHp 2’/3’-O-(N-Methyl-anthraniloyl)-guanosine-5’-[(β,γ)-imido]triphosphate)-bound TTN5, TTN5^T30N^ and TTN5^Q70L^ were prepared by incubation of nucleotide-free forms with fluorescent nucleotides (Jena Bioscience, Germany) in a molar ratio of 1 to 1.2. The solution was purified from excess amount of mdGDP and mGppNHp by using prepacked gel-filtration NAP-5 Columns (Cytiva, Germany) to remove unbound nucleotides. Protein and nucleotide concentration were determined using the Bradford reagent (Sigma Aldrich, Germany) and HPLC, respectively.

All kinetic fluorescence measurements including nucleotide association and dissociation reactions were monitored on a stopped-flow instrument system SF-61, HiTech Scientific (TgK Scientific Limited, UK) and SX20 MV (Applied Photophysics, UK) at 25°C using nucleotide exchange buffer (10 mM K_2_HPO_4_/KH_2_PO_4_, pH 7.4, 5 mM MgCl_2_, 3 mM DTT, 30 mM Tris/HCl, pH 7.5) (Eberth and Ahmadian 2009). Fluorescence was detected at 366 nm excitation and 450 nm emission using 408 nm cut-off filter for mant-nucleotides (Hemsath and Ahmadian 2005).

To determine the intrinsic nucleotide exchange rate, k_off_, 0.2 µM mdGDP- and mGppNHp-bound proteins were combined with a 200-fold molar excess of 40 µM non-fluorescent GDP in two different set of experiments, respectively. The decay of the fluorescence intensity representing mdGDP and mGppNHp dissociation and replacement by non-fluorescent nucleotide were recorded over time (Supplementary Figure S2G). Moreover, to determine the nucleotide association rate, k_on_, of mdGDP and mGppNHp to the nucleotide-free GTPase, 0.2 µM fluorescent nucleotides were mixed with different concentrations of nucleotide-free TTN5 variants. The increase in the fluorescent intensity was obtained by the conformational change of fluorescent nucleotides after binding to the proteins (Supplementary Figure S2H).

The data provided by stopped-flow were applied to obtain the observed rate constants. Dissociation rate constants or nucleotide exchange rates (k_off_ in s^-1^) and pseudo-first-order rate constants or observed rate constants (k_obs_ in s^-1^) at the different concentrations of the protein were obtained by non-linear curve fitting using Origin software (version 2021b). The slopes obtained from plotting k_obs_ against respective concentrations of proteins were used as the second-order association rate constants (k_on_ in µM^-1^s^-1^). The equilibrium constant of dissociation (K_d_ in µM) was calculated from the ratio of k_off_/k_on_. In order to investigate the intrinsic GTP-hydrolysis rate of TTN5 variants, the HPLC method is used as described (Eberth and Ahmadian 2009). As an accurate strategy, HPLC provides the nucleotide contents over time. The GTPase reaction rates were determined by mixing 100 μM nucleotide-free GTPase and 100 μM GTP at 25°C in a standard buffer without GDP. The GTP contents were measured at different times and the data were fitted with Origin software to get the observed rate constant.

### Nicotiana benthamiana leaf infiltration

*N. benthamiana* leaf infiltration was performed with the Agrobacterium (*Agrobacterium radiobacter*) strain C58 (GV3101) carrying the respective constructs for confocal microscopy. Agrobacteria cultures were grown overnight at 28°C, centrifuged for 5 min at 4°C at 5000*g*, resuspended in infiltration solution (5 % sucrose, a pinch of glucose, 0.01 % Silwet Gold, 150 µM Acetosyringone) and incubated for 1 hour at room temperature. Bacterial suspension was set to an OD_600_ = 0.4 and infiltrated into the abaxial side of *N. benthamiana* leaves.

### Subcellular localization of fluorescent protein fusions

Cloning of YFP-tagged TTN5 constructs is described in the paragraph ‘Arabidopsis plant material and growth conditions’. Localization studies were carried out by laser-scanning confocal microscopy (LSM 780 or LSM880, Zeiss) with a 40x C-Apochromat water immersion objective. YFP constructs and Alexa Fluor 488 stainings were excited at 488 nm and detected at 491-560 nm. mCherry, Alexa 555 or FM4-64 fluorescence was excited at 561 nm and detected at 570-633 nm.

Wortmannin (10 µM, Sigma-Aldrich), BFA (36 µM Sigma-Aldrich) and plasma membrane dyes FM4-64 (165 µM, ThermoFisher Scientific) were infiltrated into *N. benthamiana* leaves. FM4-64 was detected after 5 min incubation. Wortmannin and BFA were incubated for 25 min before checking the treatment effect. Plasmolysis was induced by incubating leaf discs in 1 M mannitol solution for 15 min. Signal intensities were increased for better visibility.

### Whole-mount Immunostaining

Whole-mount immunostaining by immunofluorescence was performed according to the protocol described by (Pasternak et al. 2015). Briefly, Arabidopsis seedlings were grown in the standard condition in Hoagland media for 4-6 days. Methanol or Formaldehyde (4 %) was used to fix the seedlings. The seedlings were transferred to a glass slide and resuspended in 1x microtubule-stabilizing buffer (MTSB). Seedlings were digested with 2 % Driselase dissolved in 1x MTSB at 37°C for 40 mins. Following digestion, permeabilization step was performed by treating the seedlings with permeabilization buffer (3 % IGEPAL C630, 10 % dimethylsulfoxide (DMSO) in 1x MTSB buffer) at 37°C for 20 mins. Then blocking was performed with a buffer consisting of 5 % BSA for 30 min at room temperature. They were incubated overnight with different primary antibodies (detailed information are listed below). After two washes with 1x MTSB, seedlings were incubated with a respective Alexa Fluor secondary antibody for 2 hours at 37°C. After five steps of washing with 1x PBS, coverslips were mounted on slides with the antifade reagent (Prolong glass Antifade Mountant with NucBlue Stain, Invitrogen). Fluorescence microscopy was conducted as described in the previous section.

Immunodetection was conducted with following antibody combinations: HA detection was performed using α-HA antibody (1:100 dilution, rabbit Abcam ab9110 or chicken AGRISERA, AS204463) followed by Alexa Fluor 488 or Alexa Fluor 555-labeled secondary antibodies (1:200 α-rabbit, Thermo Scientific, A32731 and AGRISERA, AS204463). ATPase (PM marker) was detected using primary chicken α-AHA antibody (1:200 Agrisera, AS132671), while ARF1 (Golgi and TGN marker) was detected using primary chicken α-ARF1 antibody (1:200 Agrisera, AS08325), both in combination with Alexa Fluor™ Plus 555-labeled secondary antibody (1:500 α-Chicken for ATPase, Thermo Scientific A32932).

For obtention of BFA bodies, seedlings were first treated with BFA (72 µM, Sigma-Aldrich) and fixable plasma membrane dye FM4-64 FX (10 mM, Thermo Scientific, F34653) for 1 hour, before formaldehyde fixation.

### Immunoblot detection

Total protein extraction from Arabidopsis plants grown for 6 days or in the 2-week system, sample separation on SDS-PAGE and immunodetection were performed as previously described (Le et al. 2015). In short, plant material was grinded under liquid nitrogen and proteins were extracted with SDG buffer (62 mM Tris-HCl, pH 8.6, 2.5 % SDS, 2 % DTT, 10 % glycerol). Samples were separated on 12 % SDS-PAGE gels. Following electrophoresis, the proteins were transferred to a Protran nitrocellulose membrane (Amersham).

Membranes were blocked for 1 hour in 5 % milk-TBST solution (20 mM Tris-HCl, pH 7.4, 180 mM NaCl and 0.1 % Tween 20), followed by 1 hour antibody incubation (α-GFP, monoclonal mouse antibody, Roche, catalog no. 11814460001, 1:1000). After three washes with TBST for 10 min each, membranes were incubated in secondary antibody (α-mouse-HRP, polyclonal goat antibody, SigmaAldrich, cat. no. SAB3701159, 1:5000) for 1 hour. HA detection was performed with a directly coupled α-HA antibody (α-HA-HRP, high-affinity monoclonal rat antibody, 3F10, Roche, catalog no. 12013819001, 1:1000). Immunodetection was performed after three washes with TBST for 10 min each, using the enhanced chemiluminescence system (GE Healthcare) and the FluorChem Q System for quantitative Western blot imaging (ProteinSimple) with the AlphaView software.

### JACoP based colocalization analysis

Colocalization analysis was carried out with the ImageJ (Schneider et al. 2012) Plugin Just Another Colocalization Plugin (JACoP) (Bolte and Cordelières 2006) and a comparison of Pearson’s and Overlap coefficients and Li’s intensity correlation quotient (ICQ) was performed. Object-based analysis was done for punctate structures, adapted by (Ivanov et al. 2014). Colocalization for both channels was calculated based on the distance between geometrical centers of signals and presented as percentage. Analysis was done in three replicates each (n = 3).

### Structure prediction

TTN5 structure prediction was performed by AlphaFold (Jumper et al. 2021). The molecular graphic was edited with UCSF ChimeraX (1.2.5, (Goddard et al. 2018), developed by the Resource for Biocomputing, Visualization and Informatics at the University of California, San Francisco, with support from the National Institutes of Health R01-GM129325 and the Office of Cyber Infrastructure and Computational Biology, National Institute of Allergy and Infectious Diseases.

### *In silico* tool for gene expression analysis

RNA-seq data relies on published data and was visualized with the AtGenExpress eFP at bar.utoronto.ca/eplant (Waese et al. 2017).

### Statistical analysis

One-way ANOVA was used for statistical analysis and performed in OriginPro 2019. Fisher LSD or Tukey was chosen as a post-hoc test with p < 0.05.

## ACCESSION NUMBERS

Sequence data from this article can be found in the TAIR and GenBank data libraries under accession numbers: *ARA7* (TAIR: AT4G19640), *ARF1* (TAIR: AT1G23490), *GmMan1* (Uniprot: Q0PKY2) and *TTN5* (TAIR: AT2G18390).

## Author contributions

I.M., M.R.A., R.I., and P.B. conceived the project; I.M., A.M., S.K.S., P.C., M.R.A., R.I. and P.B. designed experiments, supervised the research and analyzed data; I.M., A.M., S.K.S., and P.C. performed experiments and analyzed data; I.M. wrote the manuscript; I.M., A.M., M.R.A., R.I., and P.B. edited the manuscript; P.B. acquired funding. R.I. and P.B. agreed to serve as the author responsible for contact and communication.

## Abbreviations

Arabidopsis: *Arabidopsis thaliana*
ARF-like / ARL: ADP-ribosylation factor-like
BFA: brefeldin A
EE: early endosomes
GAP: GTPase-activating protein
GEF: guanine nucleotide exchange factor
MVB: multivesicular body
TGN: *trans*-Golgi network
TTN5: TITAN 5

## Acknowledgements

We thank Gintaute Matthäi and Elke Wieneke for their excellent technical assistance. We are thankful to Dr. Anna Sergeeva for advice and help with whole-mount immunolocalization. We thank Dr. Ksenia Krooß for microscopy help and advice and Natalie Köhler for experimental assistance. We are thankful for the assistance from Stefanie Weidtkamp-Peters and Sebastian Hänsch, members of the Center for Advanced Imaging (CAi) at Heinrich Heine University. We would like to sincerely thank Dr. Madhumita Narasimhan for her help with immunolocalization experiments. We are greatful to Dr. Alexander Hertle for many fruitful discussions and advices on immunolocalization and fluorescence microscopy. RFP-ARA7 clones were a gift from Dr. Thierry Gaude.

This work was supported by the Deutsche Forschungsgemeinschaft (DFG, German Research Foundation) Project no. 267205415–SFB 1208, project B05 to P.B.; Ba1610/7-2 to P.B.; Germany’s Excellence Strategy – EXC-2048/1 – project ID 390686111. Support was provided by DFG AH 92/8-3 to A.M. and M.R.A.. Funding for instrumentation: Zeiss LSM780 + 4channel FLIM extension (Picoquant): DFG-INST 208/551-1 FUGG and Zeiss LSM 880 Airyscan Fast DFG-INST 208/746-1 FUGG.

## Supplementary figures

**Supplementary Figure S1.**
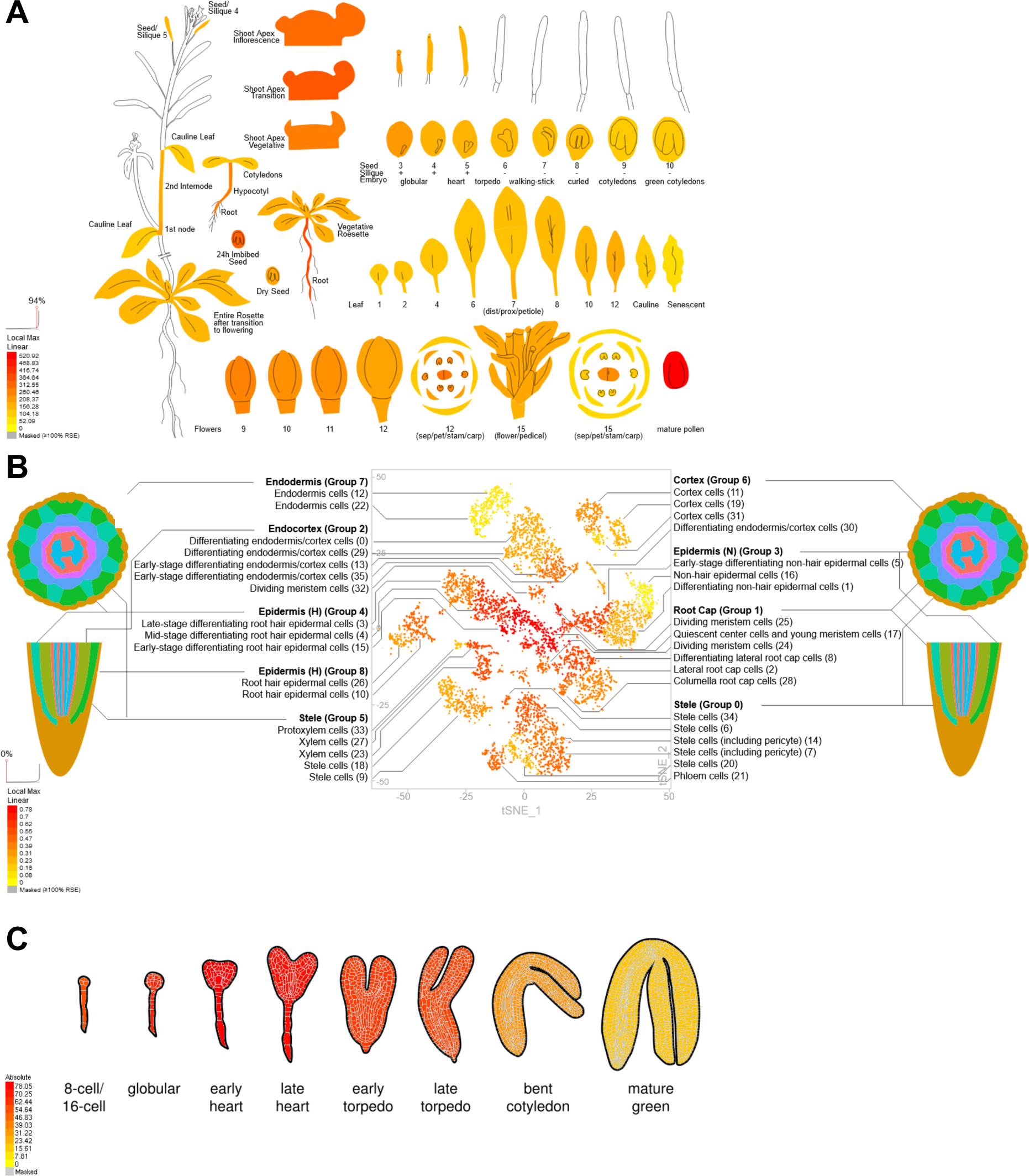
Visualization of *TTN5* gene expression levels during plant development based on transcriptome data. Expression levels in (A), different types of aerial organs at different developmental stages; from left to right and bottom to top are represented different seed and plant growth stages, flower development stages, different leaves, vegetative to inflorescence shoot apex, embryo and silique development stages; (B), seedling root tissues based on single cell analysis represented in form of a uniform manifold approximation and projection plot; (C), successive stages of embryo development. As shown in (A) to (C), *TTN5* is ubiquitously expressed in these different plant organs and tissues. In particular, it should be noted that *TTN5* transcripts were detectable in the epidermis cell layer of roots that we used for localization of tagged TTN5 protein in this study. In accordance with the embryo-lethal phenotype, the ubiquitous expression of *TTN5* highlights its importance for plant growth. Original data were derived from (Nakabayashi et al. 2005, Schmid et al. 2005) (A); (Ryu et al. 2019) (B); (Waese et al. 2017) (C). Gene expression levels are indicated by local maximum color code, ranging from the minimum (no expression) in yellow to the maximum (highest expression) in red.

**Supplementary Figure S2.**
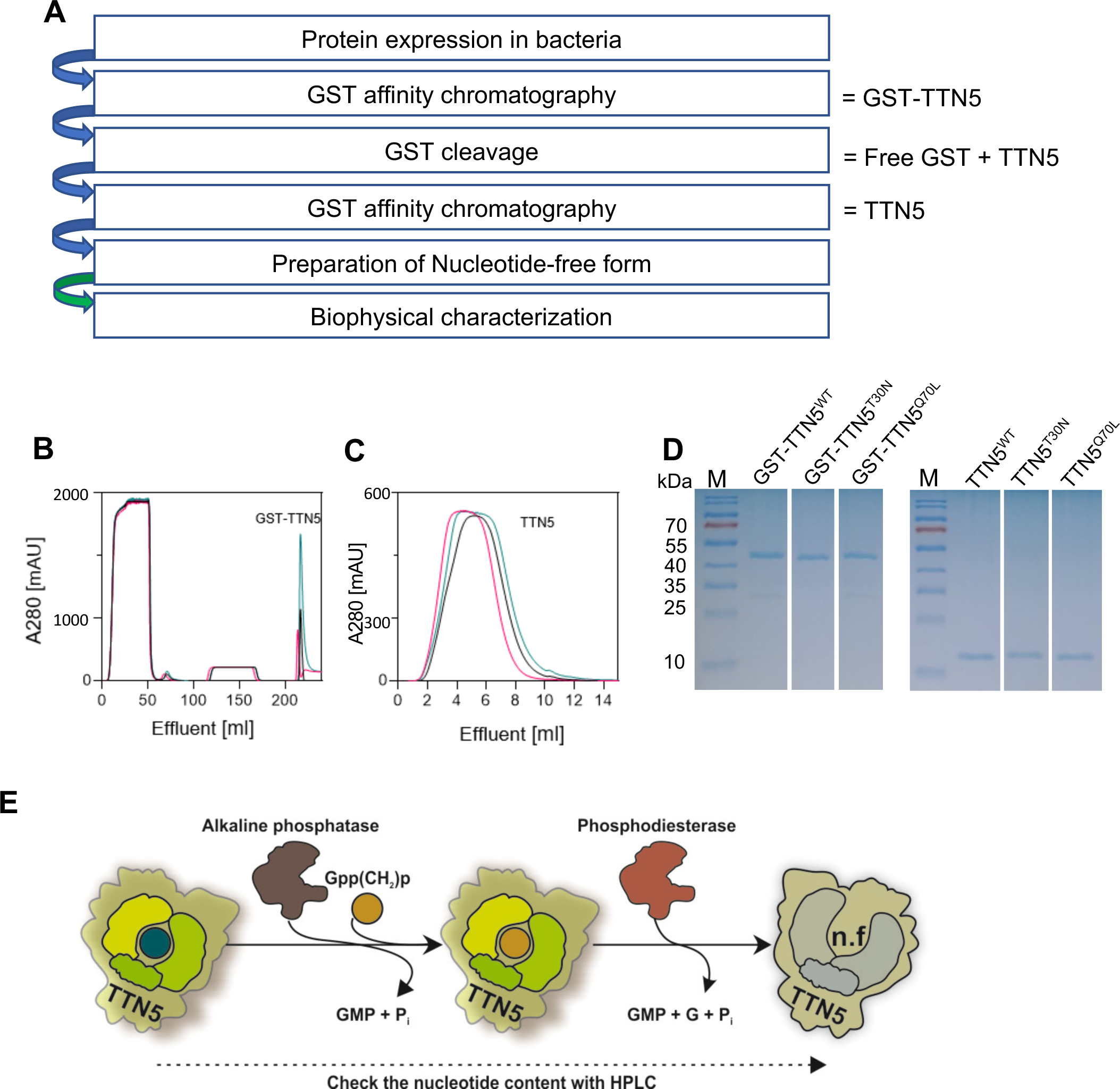
Heterologous expression and purification of TTN5 protein as well as preparation of its nucleotide-free form. (A), Overview of protein purification and preparation of nucleotide-free form of TTN5 in the presence of excess GDP. (B-C), The chromatograms represent the GST-affinity chromatography to obtain GST-TTN5 and TTN5 variants, before and after GST-cleavage by thrombin. (D), Coomassie Blue SDS-PAGE of GST-TTN5 (left panel; 46.5 kDa) and TTN5 after GST-cleavage by thrombin and a second GST-affinity chromatography (right panel; 21 kDa). (E), Schematic illustration of nucleotide-free TTN5 preparation. In the first step, the purified GDP-bound TTN5 is incubated with alkaline phosphatase in the presence of 1.5-fold molar excess of Gpp(CH_2_)p, which is a non-hydrolyzable GTP analog and, unlike GDP, resistant to alkaline phosphatase. After GDP is completely degraded, phosphodiesterase is added to the reaction to degrade Gpp(CH_2_)p to GMP, G and Pi. The nucleotide content is monitored in each step by HPLC. The solution with nucleotide-free TTN5 is deep-frozen and thawed twice, aliquoted and stored at -80°C.

**Supplementary Figure S3.**
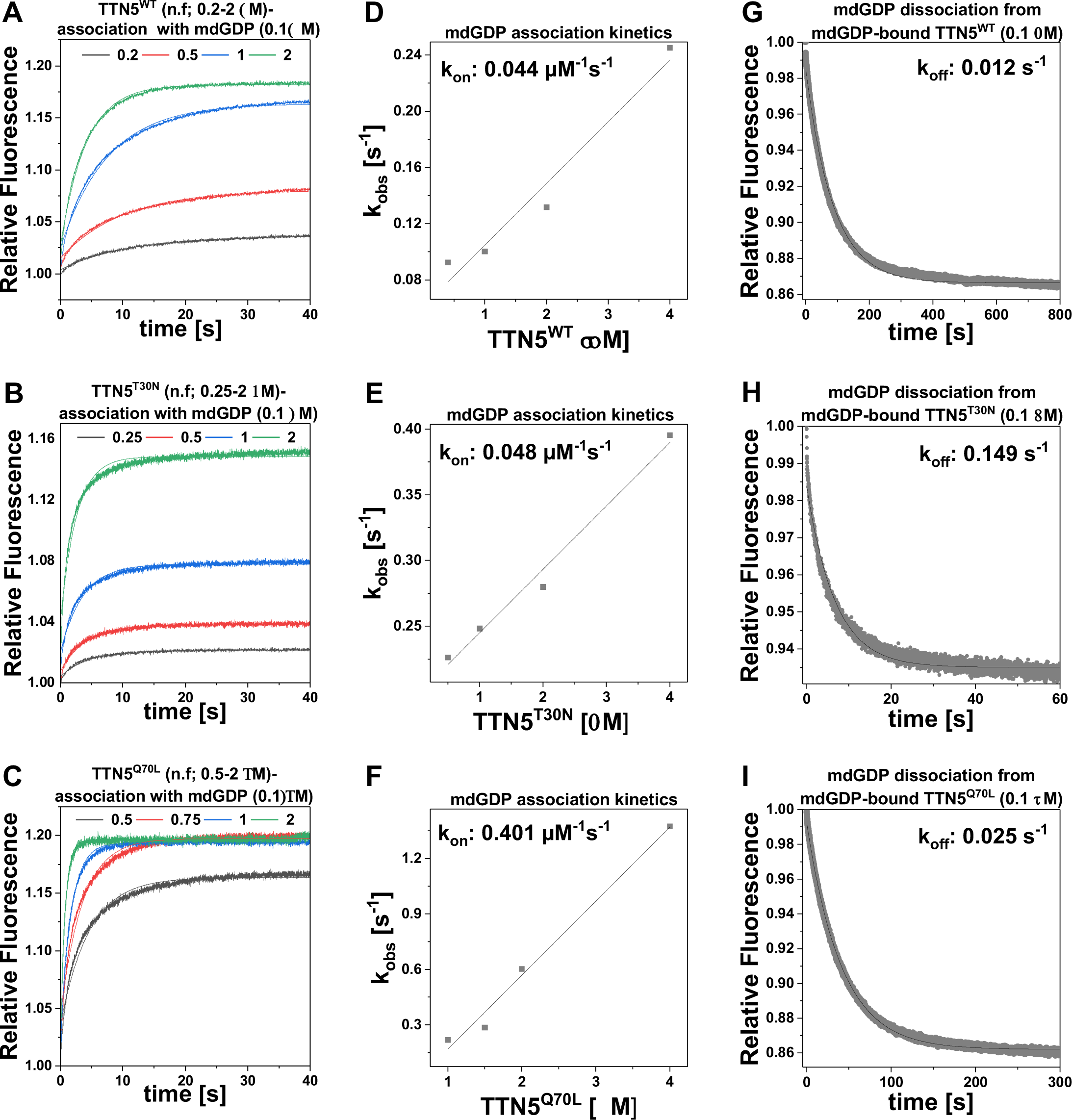
Kinetic measurements of mdGDP interaction with TTN5 proteins. (A-C), Association of mdGDP (0.1 μM) with increasing concentrations of TTN5^WT^ (A), TTN5^T30N^ (B) and TTN5^Q70L^ (C). (D-F), Association rate constant (k_on_) is determined by plotting the k_obs_ values, obtained from the exponential fits of the mdGDP association data (A-C) against the TTN5 (D), TTN5^T30N^ (E) and TTN5^Q70L^ (F) protein concentrations. (G-I), Dissociation of mdGDP from TTN5 (G), TTN5^T30N^ (H) and TTN5^Q70L^ (I) proteins (0.1 μM) is determined in the presence of excess amounts of unlabeled GDP (20 μM). The dissociation rates (k_off_) are obtained from the exponential fitting of the data by Origin software. All results are shown as bar charts in Figure 2D. The principles of the assays are illustrated in Figure 2A-C.

**Supplementary Figure S4.**
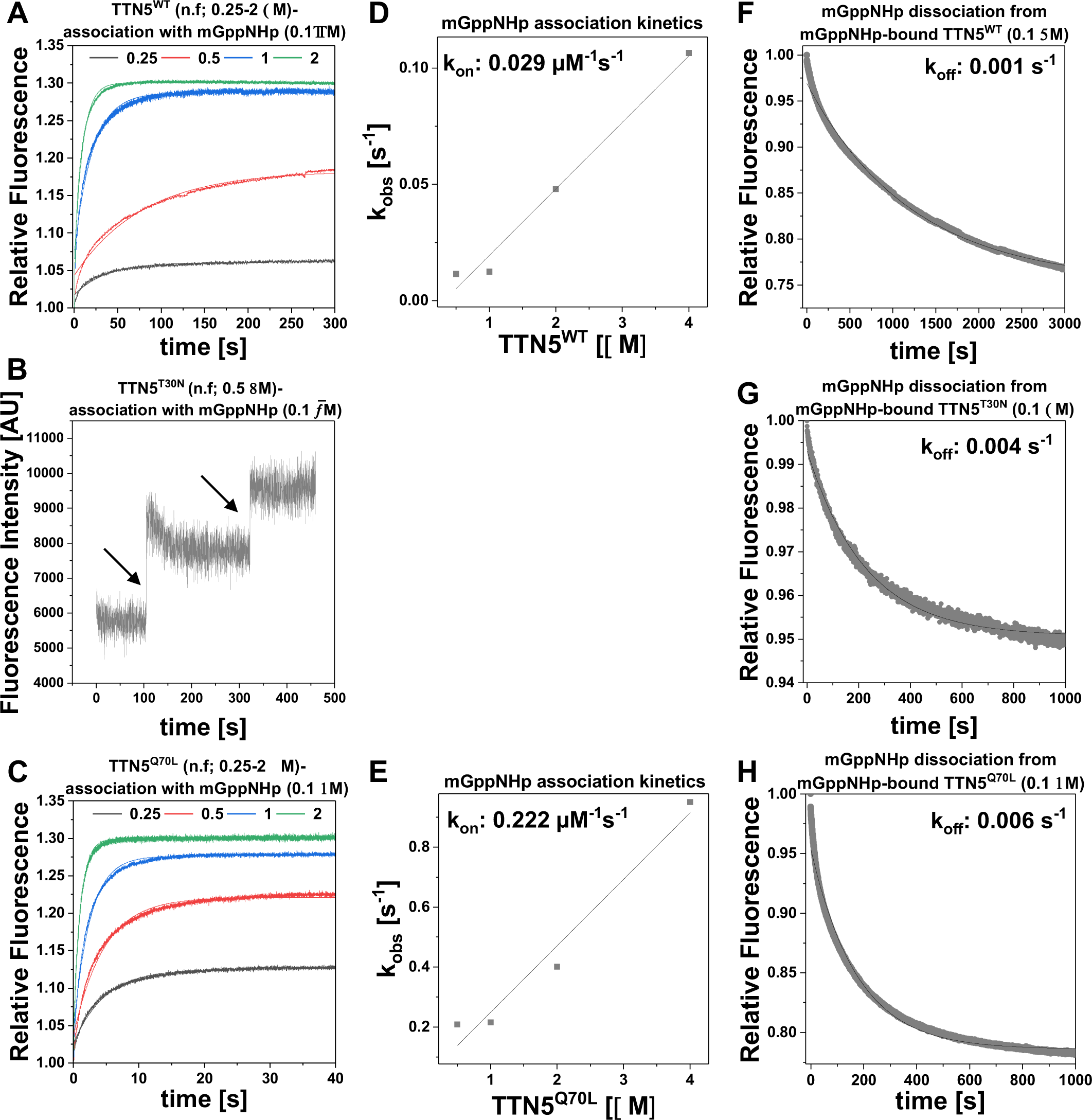
Kinetic measurements of mGppNHp interaction with TTN5 proteins. (A-E), Association kinetics. (A, C), Association of mGppNHp (0.1 μM) with increasing concentrations of TTN5^WT^ (A) and TTN5^Q70L^ (C). (B) When mixing mGppNHp with nucleotide-free TTN5^T30N^, no binding was observed under these experimental conditions. Instead, association of mGppNHp (0.1 μM) with two different titrations of 0.5 μM TTN5^T30N^ was measured, indicated by arrows, to confirm the fast binding of TTN5^T30N^ with mGppNHp by increasing fluorescence intensity, determined by fluorimeter. (D-E), Association rate constant (k_on_) is determined by plotting the k_obs_ values, obtained from the exponential fits of the mGppNHp association data (left panels) against the TTN5^WT^ (D) and TTN5^Q70L^ (E) protein concentrations. Note that TTN5^T30N^ did not show association with mGppNHp, therefore middle space is left empty. (F-H), Dissociation of mGppNHp from TTN5^WT^ (F), TTN5^T30N^ (G) and TTN5^Q70L^ (H) proteins (0.1 μM) is determined in the presence of excess amounts of unlabeled GppNHp (20 μM). The dissociation rates (k_off_) are obtained from the exponential fitting of the data by Origin software. All results are shown as bar charts in Figure 2E. The principles of the assays are illustrated in Figure 2A-C.

**Supplementary Figure S5.**
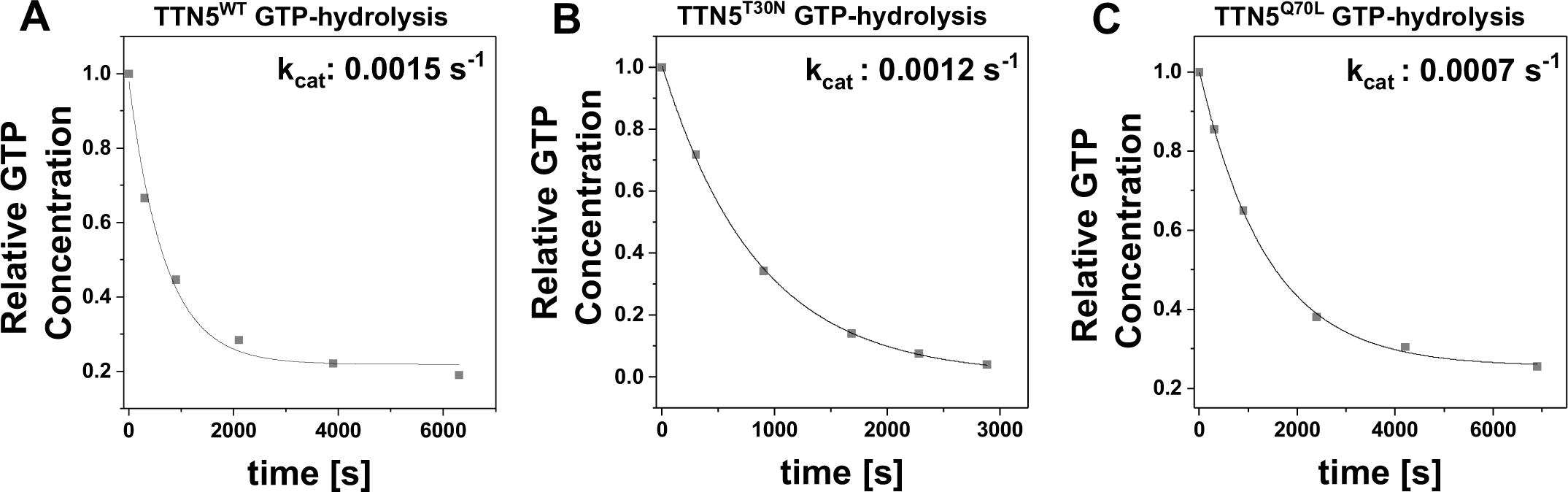
GTP hydrolysis reaction rates of the TTN5 proteins. (A-C), The GTP hydrolysis reaction rates (k_cat_) of TTN5^WT^ (A), TTN5^T30N^ (B) and TTN5^Q70L^ (C) proteins (100 µM) were measured at 25°C. Aliquots of the reaction mixture at the indicated time points were analyzed by HPLC as described in Materials and Methods. The obtained results are shown as bar charts in Figure 2G. The principle of the assay is illustrated in Figure 2F.

**Supplementary Figure S6.**
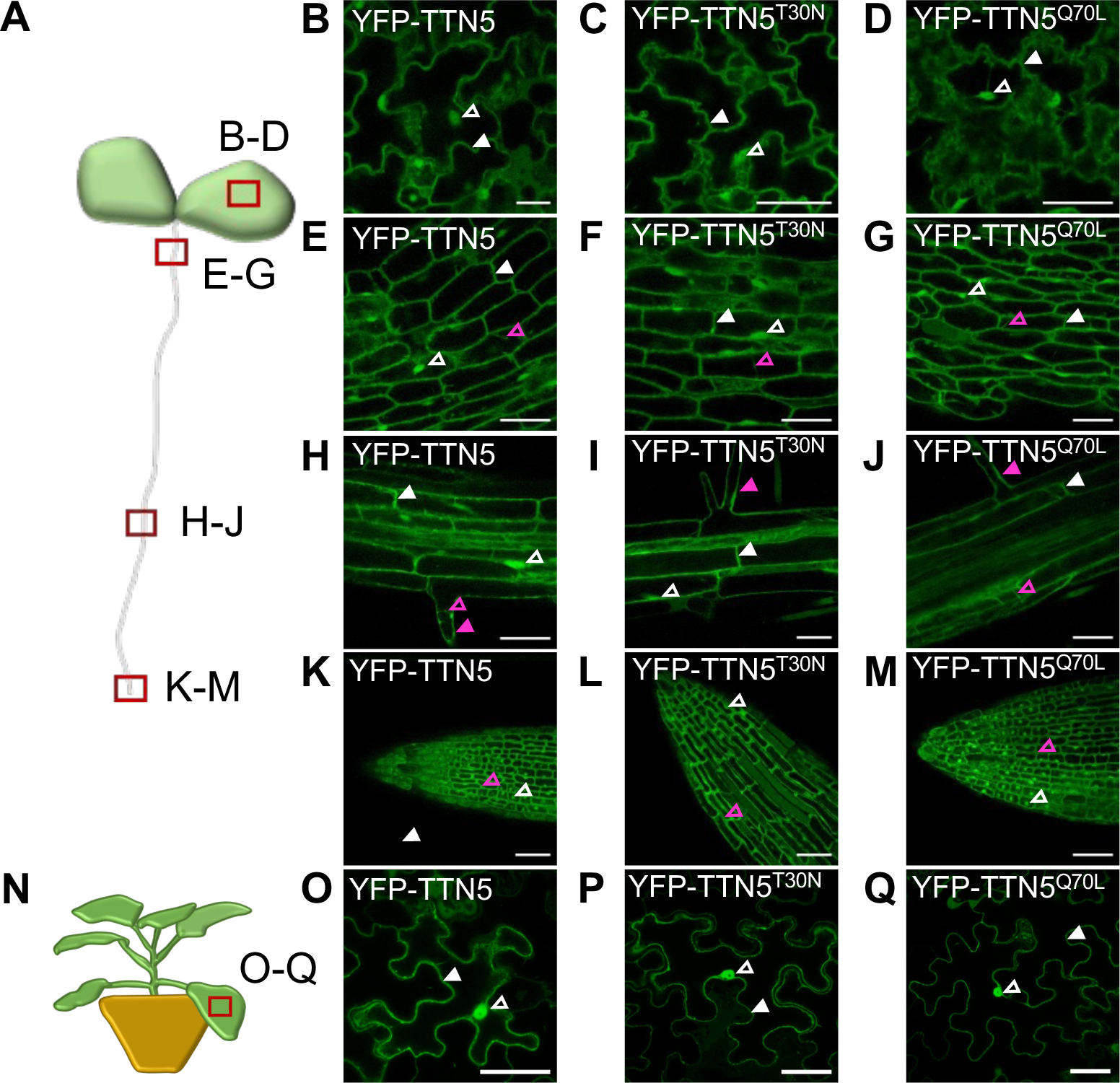
YFP fluorescence signal localization in YFP-TTN5 and -TTN5 mutant Arabidopsis seedlings. Microscopic YFP fluorescence observations were made in a plane across the centers of most cells. (A), Schematic representation of an Arabidopsis seedling. Images were taken at four different positions of the seedlings and imaged areas are indicated by a red rectangle. (B-M), Fluorescent YFP signals in Arabidopsis seedlings *via* fluorescent confocal microscopy. (B–D), YFP signal localization was observed in the epidermis of cotyledons. Fluorescence signals were present in nucleus (indicated by empty white arrowhead) and cytoplasm and at or in close proximity to the plasma membrane (indicated by filled white arrowhead) in the epidermis of cotyledons of YFP-TTN5, YFP-TTN5^T30N^ and YFP-TTN5^Q70L^ seedlings. (E-G), Fluorescence localization in the hypocotyls showed clear presence in the cytoplasm (indicated by empty magenta arrowhead), next to nuclear and plasma membrane-related localization. (H-J), Fluorescence signals were present in nuclei, cytoplasm, at or close to the plasma membrane in the root hair zone and in root hairs (indicated by filled magenta arrowhead) with clear cytoplasmic localization of YFP-TTN5, YFP-TTN5^T30N^ and YFP-TTN5^Q70L^ seedlings. (K-M), Fluorescence signal localization at the root tip showed clear expression in nuclei and cytoplasm, visible due to the smaller size of the vacuoles. (N), Schematic representation of a *N. benthamiana* plant, used for leaf infiltration for transient expression of YFP-TTN5, YFP-TTN5^T30N^ and YFP-TTN5^Q70L^. Imaged areas are indicated by a red rectangle. (O-Q), Fluorescent signals in *N. benthamiana* leaf epidermal cells *via* fluorescent confocal microscopy. Signals were present in nucleus and cytoplasm and at or in close proximity to the plasma membrane. Scale bar 50 μm.

**Supplementary Figure S7.**
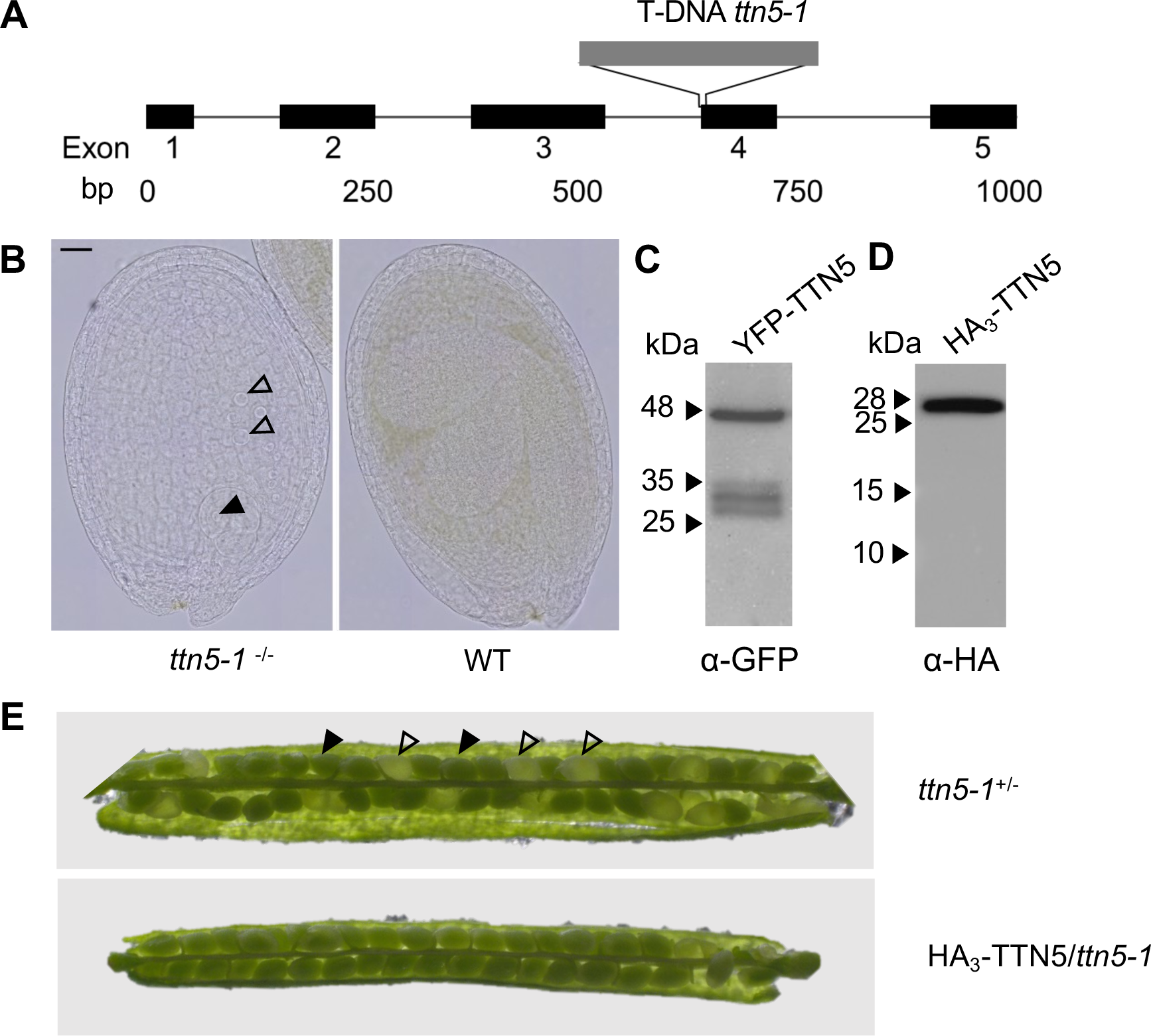
Phenotypes of *ttn5-1* T-DNA insertion line and root length of HA_3_-TTN5 lines. (A), Schematic representation of the *TTN5* exon-intron structure and T-DNA insertion in the *ttn5-1* allele. The numbers below the exon numbers indicate base pairs (bp). (B), Early embryo arrest phenotype of a homozygous *ttn5-1^-/-^* seed on the left in comparison with a wild type (WT) seed of the same silique on the right; arrowheads, indicate enlarged nuclei in the arrested embryo (filled) and endosperm (empty). Early embryo arrest phenotypes were previously shown by (Mayer et al. 1999, McElver et al. 2000). Scale bar 1 cm. (C), Western blot control of YFP-TTN5 protein expressed in stable transgenic plant lines. α-GFP primary plus secondary antibody treatment detected a strong band at the correct size of 48 kDa for YFP-TTN5 and three additional smaller sized weak bands 26-35 kDa. Plants were grown for 6 days. (D), Western blot control of HA_3_-TTN5 protein expressed in stable transgenic plant lines. One single band was detected by α-HA-HRP antibody treatment at the correct size of 28 kDa for HA_3_-TTN5 protein. Plants were grown in the two-week system. (E), Siliques of heterozygous *ttn5-1*^+/-^ plants, containing ca. 25 % white seeds with arrested embryos (empty arrowheads) and ca. 75 % regular green seeds (filled arrowheads). HA_3_-TTN5 line crossed to *ttn5-1^+/-^* was able to rescue embryo lethal seed phenotype.

**Supplementary Figure S8.**
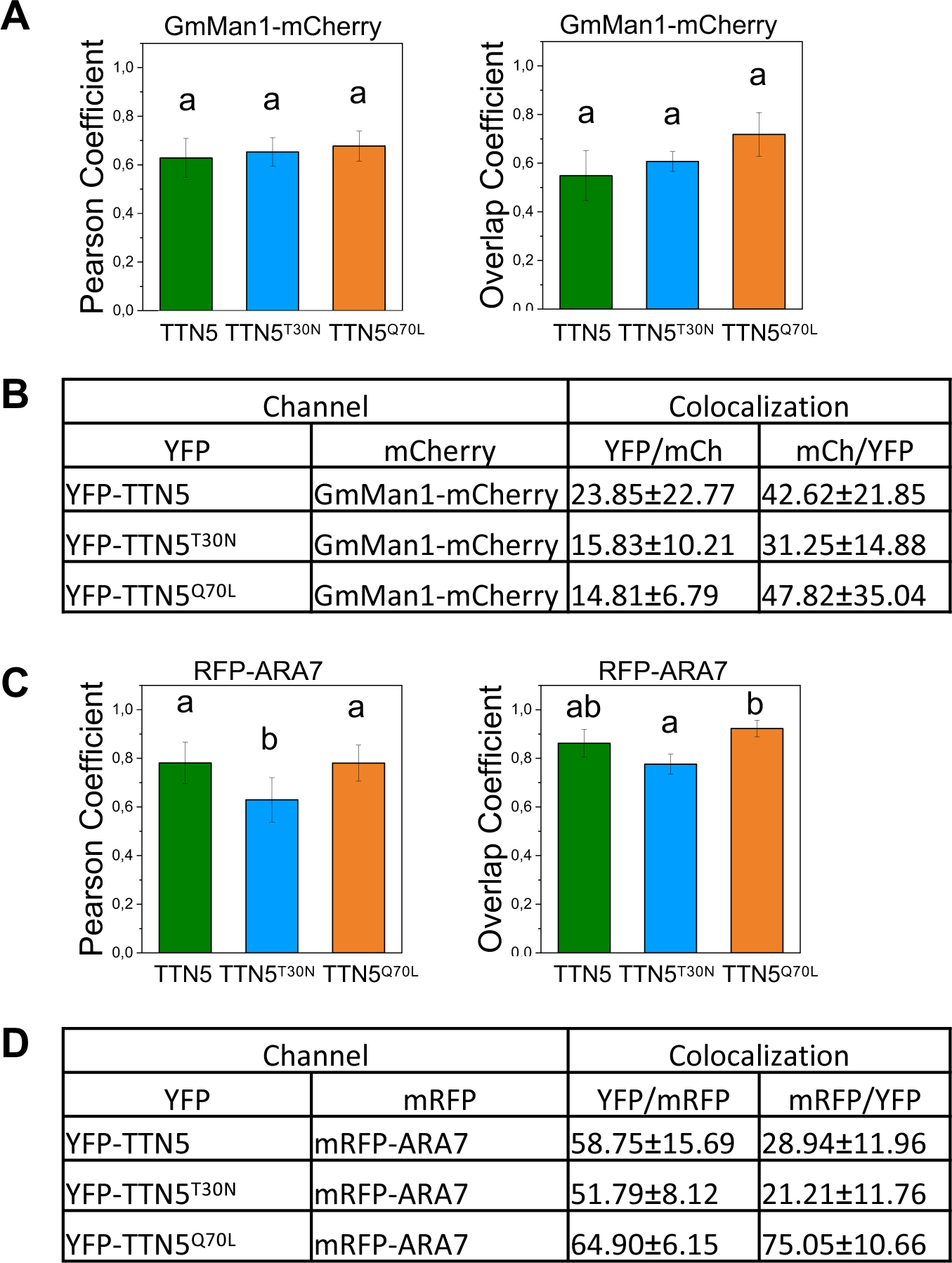
Colocalization analysis of YFP fluorescent signal with GmMan1-mCherry and mRFP-ARA7 signals in YFP-TTN5, YFP-TTN5^T30N^ and YFP-TTN5^Q70L^ seedlings. Colocalization analyses were conducted with specific markers using ImageJ (Schneider et al. 2012). (A), JACoP-based colocalization analysis (Bolte and Cordelières 2006). Comparison of Pearson’s and Overlap coefficients for *cis*-Golgi-located GmMan1-mCherry with the YFP fluorescence signals of three YFP-TTN5 seedlings in vesicle-like structures. Fluorescent signals colocalized similarly with the *cis*-Golgi marker in YFP-TTN5, YFP-TTN5^T30N^ and YFP-TTN5^Q70L^ seedlings. (B), Object-based analysis was performed for vesicle-like structures based on distance between geometrical centers of signals. YFP fluorescent signal-positive structures overlapped most with GmMan1-mCherry-positive Golgi stacks in YFP-TTN5 compared to YFP-TTN5^T30N^ and YFP-TTN5^Q70L^ seedlings. GmMan1-mCherry-positive Golgi stacks overlapped most with YFP-TTN5^Q70L^-positive structures followed by YFP-TTN5 and YFP-TTN5^T30N^ signals. (C), JACoP-based colocalization analysis (Bolte and Cordelières 2006). Comparison of Pearson’s and Overlap coefficients for TGN/MVB-located mRFP-ARA7 with YFP fluorescence signals of YFP-TTN5, YFP-TTN5^T30N^ and YFP-TTN5^Q70L^ seedlings in vesicle-like structures. YFP fluorescence signals in YFP-TTN5 and YFP-TTN5^Q70L^ seedlings colocalized similarly with mRFP-ARA7, whereas YFP signals tended to colocalize less in YFP-TTN5^T30N^ seedlings. (D), Object-based analysis was performed for vesicle-like structures based on distance between geometrical centers of signals. More than half of signals corresponding to all YFP-TTN5-positive structures overlapped with mRFP-ARA7-positive structures, in the order YFP-TTN5^T30N^, YFP-TTN5 and best YFP-TTN5^Q70L^. mRFP-ARA7-positive structures overlapped most with YFP fluorescence signals in YFP-TTN5^Q70L^ seedlings, while signals were reduced for YFP-TTN5 and YFP-TTN5^T30N^ seedlings by ca. 3.5-fold and 2.5-fold, respectively. Analyses were conducted in three replicates each (n = 3). One-way ANOVA with Fisher-LSD post-hoc test was performed. Different letters indicate statistical significance (p < 0.05).

**Supplementary Video Material S1. YFP fluorescence signals are present in dynamic vesicle-like structures in YFP-TTN5 seedlings.** (A-O), Time series of YFP fluorescent signals in dynamic vesicle-like structures of YFP-TTN5, YFP-TTN5^T30N^ and YFP-TTN5^Q70L^ Arabidopsis seedlings (A-L), or *N. benthamiana* leaf epidermal cells (M-O) recorded *via* fluorescence confocal microscopy. (A-C), Fluorescence signals were present in dynamic vesicle-like structures in epidermal cotyledon cells and stomata and in (D-F), in the hypocotyls, Note that mobility of fluorescence signal of YFP-TTN5^T30N^ (E) seedlings differed by a slower or aborted motion in half of the cells compared to (D) YFP-TTN5 and (F) YFP-TTN5^Q70L^. (G-I), YFP fluorescence signals were present in the dynamic vesicle-like structures in the root hair zone and in root hairs as well. (J-L), but no clear mobility or vesicle-like structures were deciphered in the root tips of YFP-TTN5, YFP-TTN5^T30N^ or YFP-TTN5^Q70L^ seedlings. (M-O), Time series of YFP fluorescence signals together with *cis*-Golgi marker GmMan1-mCherry. GmMan1-positive Golgi stacks showed movement in *N. benthamiana* epidermal cells together with YFP fluorescence upon transient transformation with YFP-TTN5 (M), YFP-TTN5^T30N^ (N) and YFP-TTN5^Q70L^ (O) constructs. GmMan1 is described with a stop-and-go directed movement mediated by the actino-myosin system (Nebenführ 1999) and similarly it might be the case for YFP-TTN5 signals based on the colocalization. Note that mobility of YFP fluorescence and GmMan1-mCherry signal in YFP-TTN5^T30N^ (N) transformed leaf discs differed by a slower or aborted motion compared to YFP-TTN5 (M) and YFP-TTN5^Q70L^ (O). Hence fluorescence signals were present in a comparable manner to Arabidopsis seedlings. Scale bar 50 μm.

